# In silico degradomics reveals disease- and endotype-specific alterations in the joint tissue landscape

**DOI:** 10.64898/2026.02.18.706378

**Authors:** Anna Hoyle, Kim S. Midwood

## Abstract

Tissues dynamically remodel extracellular matrix to maintain homeostasis, alterations in which are an early pathogenic hallmark of disease. Protein degradation, essential for tissue remodelling, is often dismissed as indiscriminate damage, despite evidence of its specificity. A major determinant of protein tissue levels and activity, matrix proteolysis also creates circulating degradation products that are emerging biomarkers, with specific collagen fragments capable of tracking disease severity. Understanding intentional matrix destruction therefore is key to understanding tissue biology. Unbiased, holistic analysis, extending our knowledge beyond ubiquitously expressed collagens, will uncover tissue– and disease-specific remodelling.

However, degradomics’ technical demands, requiring labelling and enrichment for neo-epitopes generated by cleavage events, restricts its inclusion in omics research. Here, we develop an in-silico pipeline (DegrAID) that identifies semi-tryptic peptides in unlabelled/unenriched proteomic datasets, mapping neo-epitopes within matrix domain organization and 3D structure, correlating these with known/predicted protease sites, and applies this to rare patient cohorts.

Validation with matched degradomic data showed good conservation across degraded proteins and cleavage sites. Interrogation of multiple, independent cohorts including cartilage, synovial tissue and synovial fluid from osteoarthritis (OA) or rheumatoid arthritis (RA) patients identified distinct degradomes between disease and tissue compartments. Further investigating RA heterogeneity revealed myeloid and lymphoid endotypes that display different treatment responses, have substantially different degradation patterns. Proteoglycans were more degraded in myeloid-RA, while collagens more so in lymphoid-RA, with notable exceptions, and endotype-specific fingerprints were conserved between synovial tissue and fluid. Thus, this tool provides new insights into tissue remodelling by unlocking degradomes from any proteomic dataset.

**One Sentence Summary:** Disease– and endotype– specific degradomes generated from clinical proteomics datasets, reveal distinct tissue remodeling patterns.

## INTRODUCTION

Tissues are architecturally complex systems, comprising cell populations embedded within highly specialised, site– and niche-specific extracellular matrix (ECM). The integrity of the tissue is dictated by the reciprocal relationship between these components, and positional cues that maintain homeostasis are balanced through dynamic deposition, remodelling and degradation of ECM. This remodelling is required for both the development and maintenance of tissue. It can however be altered early in disease, influencing external cellular cues and perpetuating pathogenic behaviour. Different diseases progress and interact with tissues in a distinct manner resulting in vastly different patterns of tissue remodelling (*1*). This can broadly reflect disease tropism for different tissue locations, as well as more refined differences, for example remodelling occurring during metastasis that is markedly different to that at the primary tumour site (*1*, *2*). As such altered ECM composition is a key hallmark of disease. Moreover, changes often precede physical symptoms such as pain and inflammation (*3*). However, the specifics of matrix remodelling in both health and disease, and any causal impact on loss of tissue homeostasis, are not fully understood.

Tissue remodelling is an intricate interplay of gene transcription, protein synthesis and post-translation modification, including degradation. A plethora of tools developed to investigate these aspects of the ECM highlight how networks made from more than 1000 secreted macromolecules are assembled in site-specific combinations in healthy tissues, and how these networks are altered in disease. Annotation of matrix genes into subcategories based on function (*4*), including structural ‘core’ matrix proteins (collagens, glycoproteins, proteoglycans) and those that bind to and interact with ECM (matrix-associated proteins such as regulators, affiliated proteins and secreted factors) enabled widespread classification of matrix transcription in tissues. Furthermore, MatrisomeDB curates increasingly large numbers of proteomic datasets, detailing high resolution protein content within the ECM of many different tissues, and in a wide range of pathological states (*4*). These resources underscore a new appreciation of the complexity, and dynamical nature, of the matrix. However, a key aspect, protein degradation, remains underexplored.

Matrix degradation does not merely constitute indiscriminate tissue damage, but is instead a key aspect of any healthy tissue. This is clearly evidenced by the requirement for MMP activity in healthy tissue (*5*, *6*). In homeostatic tissues, matrix is turned over far more dynamically than first considered, even changing in line with circadian rhythms (*7–9*). Physiological matrix degradation maintains a key balance over tissue levels of proteins, and is vital for creating and sustaining tissue niches that dictate cell function and influence tissue biomechanics. This process is also an important regulator of protein signalling, it can enable proteins to lose or alter their function through destruction of specific domains or regions; and is essential for the production of bioactive matrix fragments known as matrikines that play key physiological roles. Endostatin, a collagen XVIII fragment, and tumstatin, a collagen IV fragment for example both act as endogenous angiogenesis inhibitors (*10*, *11*). Both have effects on endothelial cells with endostatin inhibiting cell migration and tumstatin inducing apoptosis, interestingly functioning through different integrins and cell signalling pathways (*12*).

It is also well established that imbalance in matrix destruction is a hallmark of disease. Indeed, detection of circulating levels of collagens I, II and III have long been used as markers of tissue destruction and loss of organ function in joint disease (*13*). Rather than focusing solely on fragments reflecting late-stage disease, more recently, early changes in matrix turnover have been detected that are not simply a result of disease, but are also linked to cause. The identification of both beneficial and pathological roles for matrikines is beginning to shed light on how disease progression can be modulated at a mechanistic level. For example, endotrophin, released by collagen VI proteolysis and upregulated in fibrosis, has been shown to initiate proinflammatory and profibrotic responses (*14*, *15*) and endostatin, increased in response to cancer, has an anti-angiogenic function acting to inhibit the formation of blood vessels therefore slowing cancer progression (*16*, *17*). Finally, more recent biomarker investigation has demonstrated how disease development or treatment response can be reflected by changes in circulating matrix fragments in diseases including fibrosis, ulcerative colitis, RA and other arthropathies (*18–22*). Despite this, our understanding of how matrix degradation occurs in health, the impact of this on tissue homeostasis, and how this process is altered and can drive disease progression, is not yet fully understood.

Degradation can be investigated through ELISA based detection of matrix fragments using antibodies recognizing neoepitopes generated upon proteolytic cleavage (*23*, *24*). This has demonstrated the potential of monitoring tissue remodelling but has largely focused on a few selected fragments originating from highly abundant collagens, neglecting the majority of the ECM. Moreover, as collagen turnover isn’t tissue or disease specific this approach lacks precision. Degradomics can be used to study proteome-wide tissue remodelling using a specific experimental workflow whereby degraded fragments are labelled and biochemically enriched, following which LC-MS/MS is used to identify and quantify labelled fragments. This technique has been applied to the study of osteoarthritis and endocarditis (*25*, *26*). Here, specific proteases of interest have been characterised in osteoarthritic cartilage and synovial fluid and their cleavage products have been detected (*27*). Requiring additional equipment, experience and cost this approach is rarely included in multi-omics approaches.

The DegrAID pipeline presented in this study bypasses the experimental burden of this approach, instead enabling analysis of any proteomic dataset to provide new insights into tissue remodelling. Here we apply this to study changes in tissue turnover at proteome scale in chronic inflammatory and degenerative joint diseases, revealing how the degradome can resolve not only different tissue compartments and different diseases, but also different endotypes of the same disease. Moreover, we show that degradomic fingerprints from tissue are also reflected in synovial fluid. These data open the door for 1) better understanding of the functional, and pathogenic, consequences of matrix breakdown and 2) identification of specific fragments capable of disease stratification and predicting treatment response. By generating valuable insights into disease specific tissue remodelling and identifying endotype-specific matrisome fragments, this analysis pipeline demonstrates strong potential for wider application in the analysis of other datasets and diseases.

## RESULTS

### Conservation of semi-tryptic and labelled degradomic analysis

To date, degradomics has largely been investigated using non-‘standard’ sample preparation methods which label degraded fragments at their exposed termini and then enrich for them prior to LC-MS/MS analysis, here termed ‘experimental degradomics’ (Figure 1A). However, a new, analytical method detecting semi-tryptic peptides is emerging as a powerful tool to search for degraded fragments directly using data generated from ‘standard’ LC-MS/MS (*28*). Whilst this may not achieve the same depth, it enables vast amounts of degradomic investigation using ‘standard’ proteomic datasets, including from rare, unique or difficult to obtain tissue samples from patient cohorts. During sample preparation for LC-MS/MS, proteins are digested using trypsin which cleaves specifically after lysine or arginine residues, meaning that tryptic peptides are highly predictable. Semi-tryptic peptides are those with only one termini resulting from trypsin digestion as opposed to both, and are therefore indicators of in vivo protein degradation, arising as a result of protein cleavage at the other terminus by tissue resident proteases (Figure 1A).

**Fig. 1.**
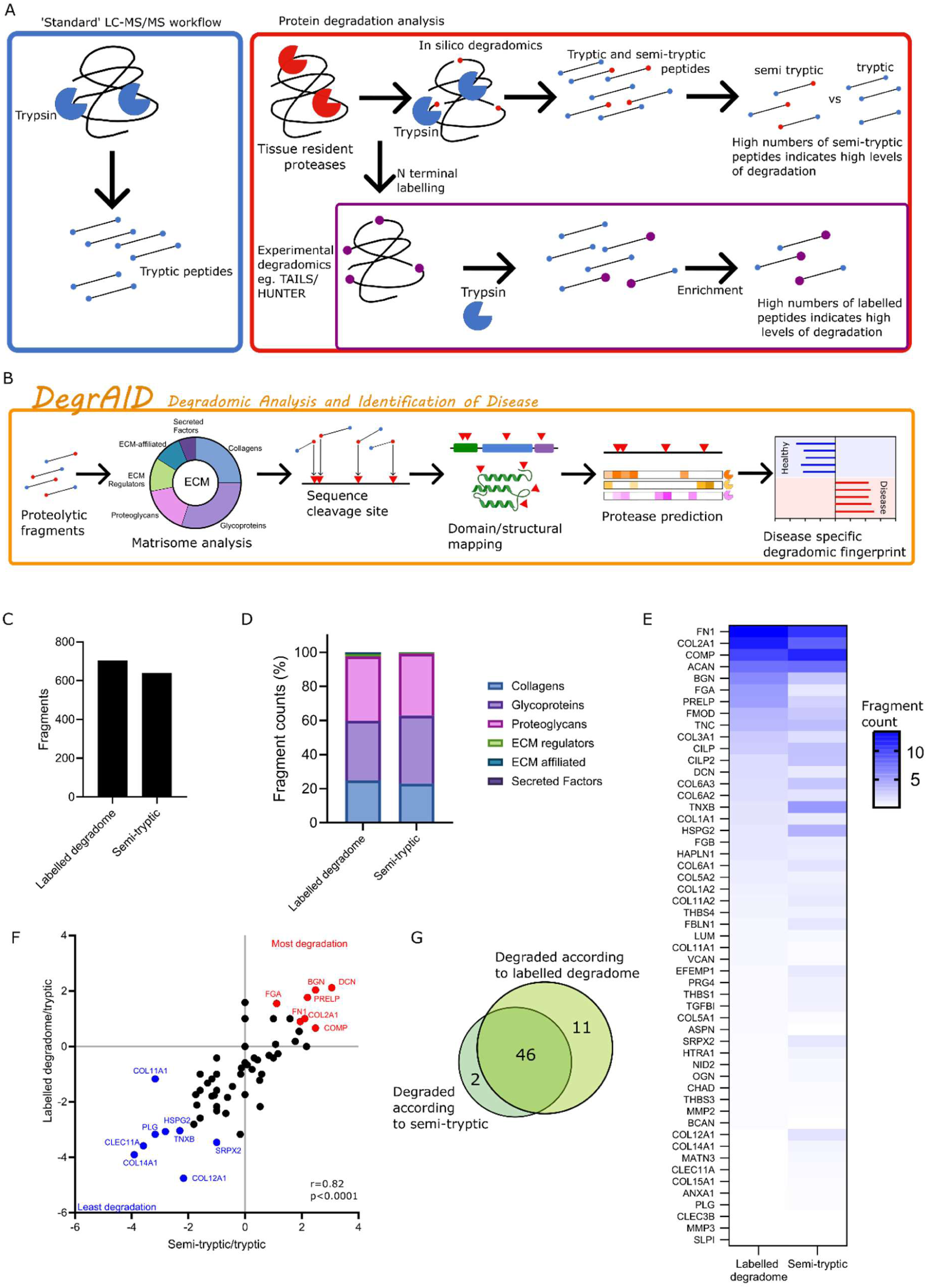
Using semi-tryptic peptides to investigate the degradome. (A) Schematic outlining the use of semi-tryptic peptides in degradome analysis and how this compares to standard proteomic and degradomic pipelines. (B) Total fragment counts resulting for labelled degradome or semi-tryptic analysis. (C) Distribution of fragments according to matrisome structure/function classifications described by Shao *et al.* (*4*). (D) Fragment counts of individual matrix proteins as detected by either labelled degradome or semi-tryptic analysis. (E) Semi-tryptic to tryptic peptide ratios (Log2) for matrix proteins detected by either labelled degradome or semi-tryptic analysis. Proteins labelled in red indicate those with the most degradation and those in blue represent the most stable proteins. Significant association was determined by Pearson correlation (r=0.82, p<0.0001). (F) Overlap between the 2 degradomic techniques described here according to analysis in (E).

Here we present DegrAID (Degradomics Analysis and Identification of Disease), a novel pipeline building on semi-tryptic peptide detection (*28*) which further: 1) compares semi-tryptic to tryptic abundances to determine relative rates of degradation, 2) conducts matrisomal analysis to enrich for matrix-focused information, 3) maps cleavage sites to specific ECM proteins, domains and structures and 4) matches fragmentation to predicted protease cleavage patterns sites, all to generate biologically relevant degradomic signatures of disease (Figure 1B). Additionally, we have validated this in-silico approach by directly comparing it to experimental degradomics on the same samples and we compared the consistency of the approach across data sets from the same tissue from different labs.

To compare in silico and experimental degradomics, we took advantage of publicly available datasets produced by the Apte lab comprising cartilage tissue from 5 patients with osteoarthritis (OA) in which matched patient samples were collected before and after enrichment for degraded fragments called Terminal Amino Isotopic labelling of substrates (TAILS) (*25*). Mining semi-tryptic peptides in samples pre-enrichment allowed the direct comparison of fragment detection and cleavage sites to those detected in matched samples using TAILS. Fragments here refers to peptides with a non-tryptic cleavage, these can either be identified using semi-tryptic analysis as part of an in-silico approach, or detected using a label following enrichment in experimental degradomics, in this case TAILS.

Fragment counts were comparable, with 862 semi-tryptic peptides detected compared to 1004 labelled peptides (Figure 1C). Tissue remodelling is largely characterised by extracellular protein turnover, reflecting this over 80% of both the semi tryptic (83%) and labelled (88%) degradome were fragments of extracellular matrix proteins. We therefore focused on degradation of these proteins. Matrisome analysis of these peptide fragments identified in OA cartilage revealed they originated from 64 and 80 matrix proteins respectively for in silico compared to the experimental approach. Matrix compositions between both methods were similar with the core matrix making up 98-99% of fragments in both approaches. Investigating this in more detail found that all matrix groups made up very similar amounts of the degradome using both approaches (Figure 1D). Glycoproteins showed the largest variation with 35% or 40% of fragments in in silico and experimental methods respectively compared to collagens (25% and 23%) and proteoglycans (38% and 36%). Further to this, fragment counts for individual proteins were compared and these counts were found to correlate significantly with proteins producing higher or lower numbers of fragments largely matching between approaches (Supplementary Figure 1A). In more detail, collagen type II (COL2A1), fibronectin (FN1) and cartilage oligomeric matrix protein (COMP) were within the top 3 most fragmented proteins across both methods and 9 of the top 10 were also conserved (Figure 1E).

Next, we assessed protein turnover detected by in silico compared to experimental methods. The number of degraded fragments (fragment counts) was compared to total tryptic peptide counts to give an indication of the overall dynamic status of a protein. After results were logged, positive values indicated more fragments were detected than tryptic peptides suggesting a high degree of degradation, referred to here as ‘actively degraded’. The reverse, more tryptic peptides than fragmented ones, indicated lower levels of turnover and were termed ‘stable’. In order to see whether the dynamic status of protein’s was conserved across the two methods, this ratio was calculated using either semi-tryptic (in-silico) or labelled (experimental) fragments compared to tryptic peptides and these two values were shown to correlate significantly (Figure 1F). In total 46 proteins were identified as the same status (actively degraded or stable) using both methods and only 2 were identified as degraded using in silico degradomics but weren’t using the experimental approach (Figure 1G). This suggests that biological conclusions made about the degradative status of individual proteins can be made using this new, in silico method. Plotting detected fragmentation onto specific protein sequences also reveals considerable overlap between both methods suggesting the specific cleavage sites are conserved across methods (Supplementary Figure 1B,C,D).

To assess the consistency of semi-tryptic peptide detection across different datasets, raw data from OA cartilage generated by the Apte lab was compared to OA cartilage data generated by the Cillero-Pastor lab (*29*). Both datasets were run through the DegrAID pipeline. Semi-tryptic fragments made up 45% and 37% of the total matrisome peptides detected from the Apte and Cillero-Pastor lab’s respectively (Figure 2A,B). The latter detected substantially more peptides in total which can be accounted for through variations in sample preparation and MS machines but the similar percentages of fragment detection suggest these fragments are broadly conserved across datasets. Degradation ratios, calculated as above, also showed significant correlation (Figure 2C). Literature supports the detection of fibromodulin (FMOD), prolargin (PRELP), decorin (DCN) and COMP as proteins identified here as being the most degraded in OA cartilage, with degradation of the latter being an established marker of OA and proteoglycan breakdown being widely reported in the disease (*30*, *31*). The overlap between datasets detecting proteins as either ‘actively degraded’ or ‘stable’ is also robust with 39 proteins matching, compared to 9 with different classifications using this technique (Figure 2D).

**Fig. 2.**
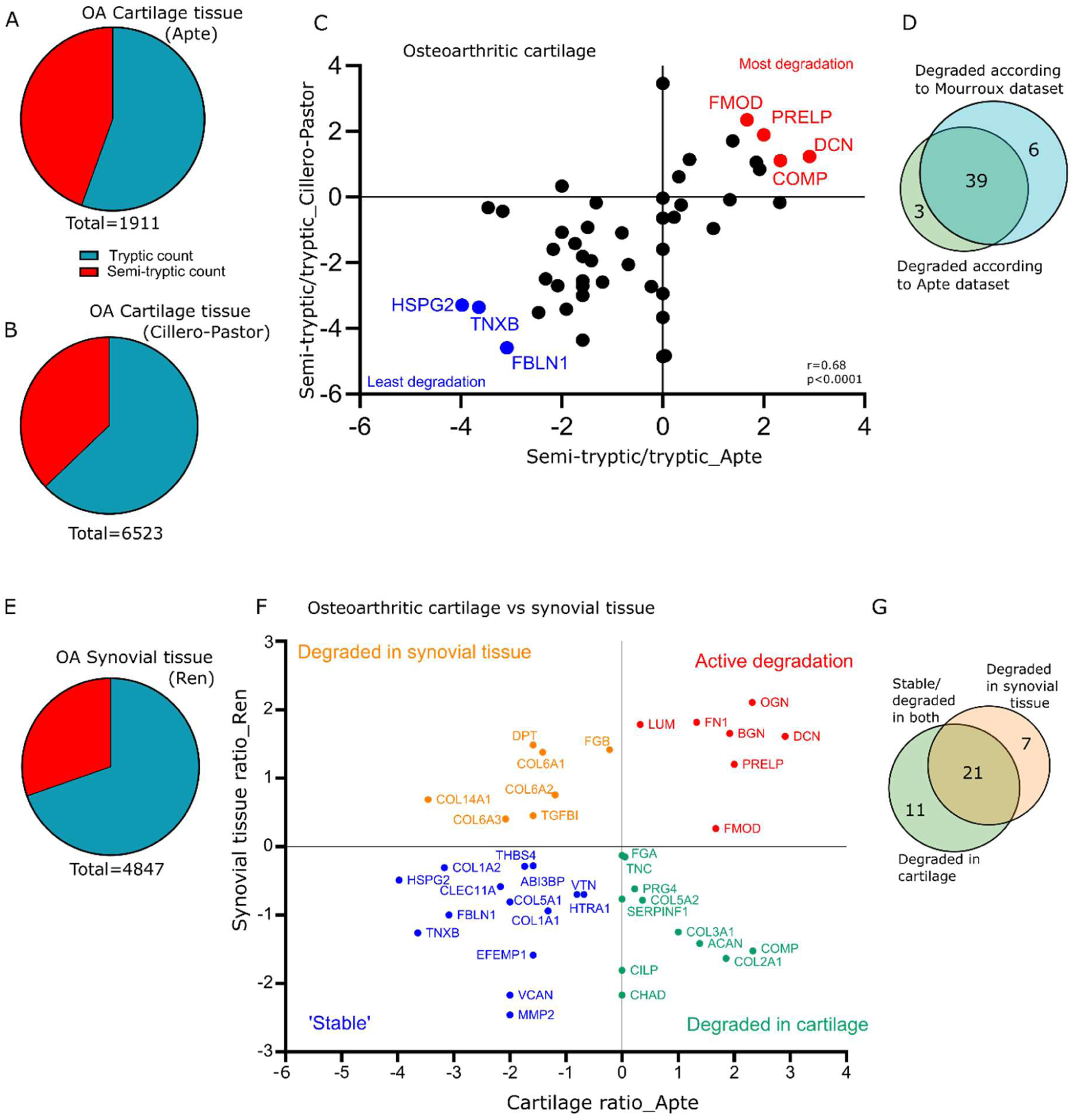
Variation in degradome between datasets and tissues. (A) Comparison of semi-tryptic (blue) and tryptic (red) peptide counts detected in OA cartilage tissue from the Apte lab (*25*). (B) As in (A) for OA cartilage tissue from the Cillero-Pastor lab (*29*). (C) Semi-tryptic to tryptic peptide ratios (Log2) for matrix proteins detected in OA cartilage tissue from either the Apte or Cillero-Pastor lab. Significant association was determined by Pearson correlation (r=0.68, p<0.0001). (D) Overlap between the different OA cartilage datasets according to analysis in (C). (E) As in (A) for OA synovial tissue from the Ren lab (*32*). (F) Semi-tryptic to tryptic peptide ratios (Log2) for matrix proteins detected in OA cartilage (Apte lab) compared with OA synovial tissue (Ren lab). (G) Overlap between the different OA tissue datasets according to analysis in (F).

Following validation of significant overlap in in silico generated degradomes from different datasets, we next examined tissue specific degradation patterns using datasets generated from cartilage or synovial tissue from individuals with OA. Semi-tryptic fragments made up 30% of the total peptidome in synovial tissue, around 10% lower than was detected in cartilage (Figure 2E). The cartilage degradome was almost solely made up of core matrix proteins whereas a quarter of the synovium degradome originated from matrix-associated proteins (Supplemental figure 1F). Relative amounts of collagen degradation were comparable but much less of the synovial degradome resulted from proteoglycan or glycoprotein degradation. Significant levels of degradation originated from ECM-regulators in synovial tissue (15%) compared to just 1% in cartilage tissue. Together, these data indicate that each tissue has a specific degradative fingerprint. In support of this, the degradative ratio between the two tissues was distinct (Figure 2F). Whilst 21 proteins displayed the same status in both tissues identifying consistent degradation or stability in a subset of proteins across cartilage and synovial tissue, 18 proteins differed between the two (Figure 2G). Collagen type VI for example has an increased degradation rate in the synovium compared to collagen type II which is more degraded in the cartilage. More widely this seems to hold true for other fibrillar compared to non-fibrillar collagens with the former more degraded in cartilage and the latter in the synovium. Interestingly, both aggrecan (ACAN) and COMP, markers of cartilage degradation in OA, showed active degradation in cartilage but relative stability in synovial tissue, suggesting this degradation is tissue specific.

Based on these data, semi-tryptic, in-silico degradomics can be used as a biologically relevant and reliable proxy for experimental alternatives, supporting the reanalysis of publicly availably proteomic datasets to probe for degradation. Consistency between datasets but variability between tissues also suggests the tissue-specificity of degradomic signatures.

### In silico degradomes distinguish OA and RA synovia, detecting high levels of RA specific collagen-VI degradation

In order to explore whether degradomic analysis can offer new insights into changes in tissues occurring as a result of disease, and if so, whether changes occur in a disease-specific manner we reanalysed a dataset produced by Ren et al., (*32*). This dataset includes Tandem Mass Tag (TMT) labelled synovial biopsies from RA (n=10) and OA (n=12) patients pooled into 6 proteomic samples. TMT labelling is a quantification method which involves chemically tagging the N-termini of peptides to allow the calculation of relative protein and peptide abundances between samples. When analysed using the DegrAID workflow described here, a total of 1472 matrisome fragments were detected. Most of the fragments originated from glycoproteins (485) and collagens (419) but a substantial number also came from proteoglycans (210) and matrix regulators (222) (Figure 3A). The proteins with the largest number of degraded fragments were largely core matrix proteins; this could reflect their higher abundance and/or their susceptibility to degradation. Different collagens and glycoproteins showed large variation in their amount of fragmentation with fibronectin and collagen type VI alpha 3 (COL6A3) exhibiting the highest fragment count, both having over 100 fragments whilst collagen IV and laminins were less degraded with under 10 fragments detected (Supplemental Figure 2A-C). A large proportion of proteoglycans displayed heavy fragmentation with perlecan (HSPG2), lumican (LUM), mimecan (OGN), DCN, PRELP and biglycan (BGN) all featuring in the top 20 most fragmented proteins suggesting that proteoglycans are heavily degraded during arthritic pathologies (Supplemental Figure 2B).

**Fig. 3.**
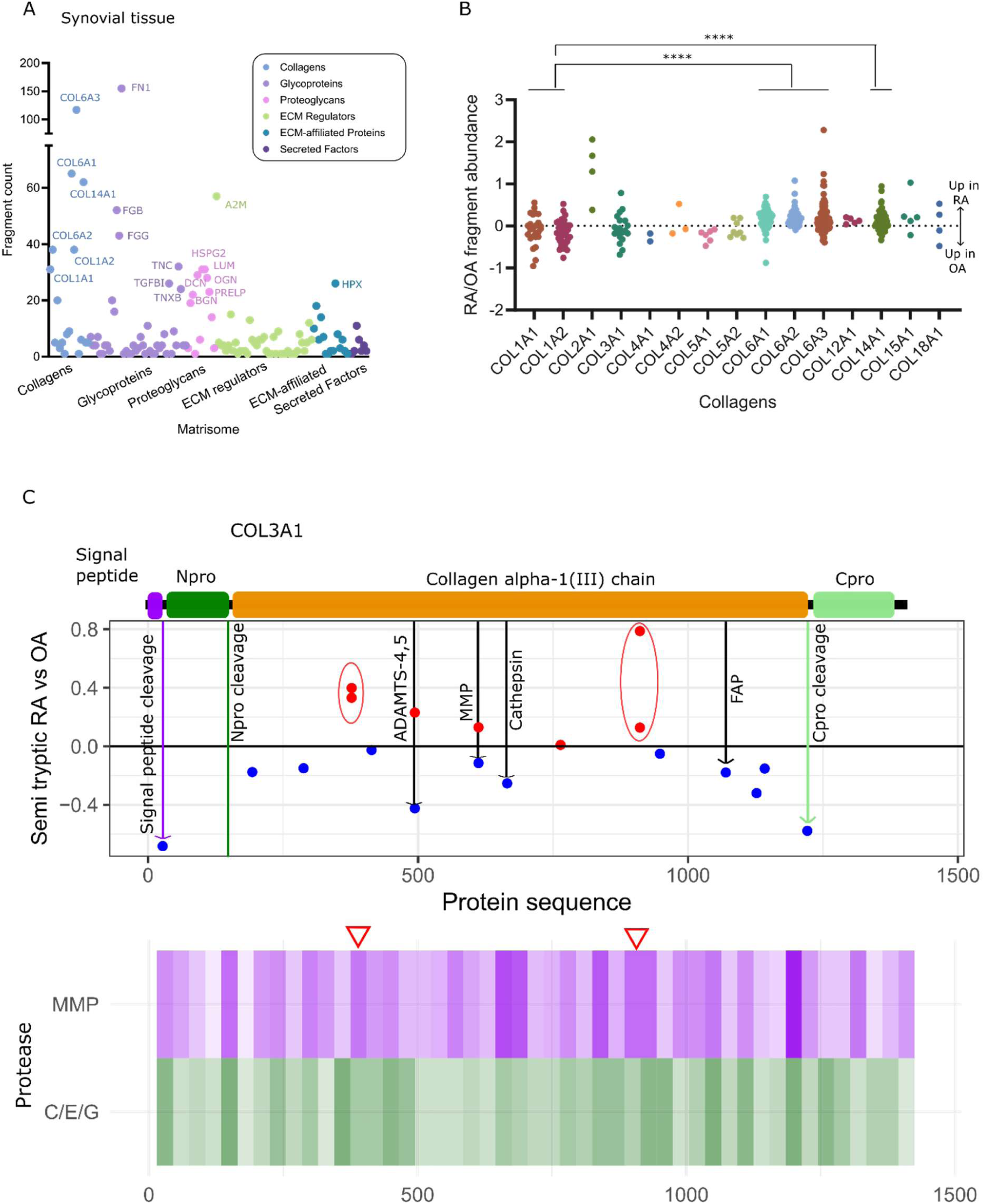
Comparison of Osteo– and Rheumatoid arthritis degradomes. (A) Total fragment counts for ECM proteins detected in synovial tissue from OA/RA patients coloured by their matrisome group (*4*). The 20 proteins with the most fragments were labelled. (B) Collagen fragment abundance (Log2) in RA compared to OA patients. Each dot relates to a different semi-tryptic peptide detected. (C) Relative abundances of detected cleavage sites in RA and OA plotted along the sequence of Collagen alpha-1(III) with a structural cartoon above. Fragments with increased in RA or OA are coloured in red or blue respectively. Vertical lines indicate cleavage sites what were either structurally indicated (coloured lines) or matched to previously reported fragments (black). The matched protease map below indicates frequency of detected cleavage sites according to Manchester Proteome so MMPs in purple and cathepsins/elastase/granzyme b in green (*92*). Darker colours correlate to increased numbers of cleavage sites. Red arrows dictate sites where multiple fragments align to a specific cleavage site upregulated in RA patients.

Having assessed the degradomic coverage of these samples we next sought to investigate the disease specificity by comparing the abundances of fragments between RA and OA. Collagen-I showed a bias towards degradation in OA synovium when its fragment abundance was compared between the two diseases (Figure 2B). Collagen VI and XIV, however, showed increased degradation in RA and this difference between collagens was significant. Crucially, this wasn’t just reflective of protein levels which actually showed the opposite trend with higher collagen-VI and XIV abundance in OA along with decreased collagen-I. Disease specific degradation was also shown in glycoproteins with, for example, tenascin-X (TNXB) showing more degradation in RA whereas tenascin-C (TNC) was more degraded in OA (Supplemental Figure 2E).

Proteoglycans, interestingly, were largely more degraded in RA with the exception of perlecan (HSPG2) which was more degraded in OA (Supplemental Figure 2D).

Next, we mapped specific cleavage sites in matrix molecules and compared these to known cleavage sites. Plotting the detected cleavage sites by the fold change in abundance in RA compared to OA demonstrated not only that this method can detect previously identified cleavage sites but also novel cleavage sites. Moreover, we also detected disease specific differences in fragmentation – both in the abundance and in the pattern of cleavage (Figure 3C). For example, a number of reported collagen-III alpha chain 1 (COL3A1) cleavage sites were detected including those resulting from cathepsin, metalloproteinases (MMPs), a disintegrin and metalloproteinase with thrombospondin motifs (ADAMTS) and fibroblast activation protein (FAP) driven proteolysis (*33–35*). Additionally, fragments resulting from signal peptide and C terminal pro-peptide cleavage were detected. Whilst protease fragments didn’t show a clear bias to either RA or OA, the processing cleavage products were upregulated in OA samples. This would suggest that whilst degradation of collagen-III largely occurs in both diseases, there is increased synthesis in OA. Detection of all of these cleavage products at one time is a significant advantage over alternative methods. Further, this pipeline identifies additional cleavage sites, both detected multiple times (circled, Figure 3C). Comparison with protease susceptibility predictions, revealed that these novel cleavage sites correlate with areas of high MMP susceptibility (arrows, Figure 3C).

In addition to matching to protease susceptibility and cleavage sites, fragments that we detected also correlated with domain organisation, protein structure and processing. For example, the fragment map for DCN showed clear clustering around 4 cleavage sites (Supplementary Figure 3A,B). These sites were all upregulated in RA patients. Interestingly, when mapped onto the DCN structure these 4 sites all match to loops, 2 of them sitting between beta sheets and 2 in areas of less defined structure. All of these regions would be more accessible and therefore more susceptible to proteases or damage. Further, the two most upregulated sites in RA appear to be the most exposed. Decorin fragments generally also show increased abundance in RA patients (Supplementary Figure 2B).

Cleavage sites originating from known protein processing events were detected for multiple collagens. These included, N-pro peptide cleavage in collagen-I and both signal peptide and endotrophin cleavage in COL6A3 (Figures 4A,B,E). Finally, protein domains also seem to influence cleavage site frequency. Collagen-VI triple helical domains were largely un-cleaved with fragment detection concentrated in von Willebrand factor (VWF) domains (Figures 4C,D,E). Interestingly, this avoidance of collagen triple helical domains isn’t seen in collagen-I indicating a difference between fibrillar and non-fibrillar collagens. Fewer fragments were detected for other collagens but cleavage sites within helical domains of fibrillar collagens II and V were detected (Supplementary Figure 3C-E). Collagen XV and XVIII both have known cleavage products (Restin and Endostatin respectively), prior to which the collagen proteins have protease sensitive regions which allow for their release. Fragments were detected within these regions for both collagens suggesting the presence of these products, particularly in RA (Supplemental Figure 3G,H).

**Fig. 4.**
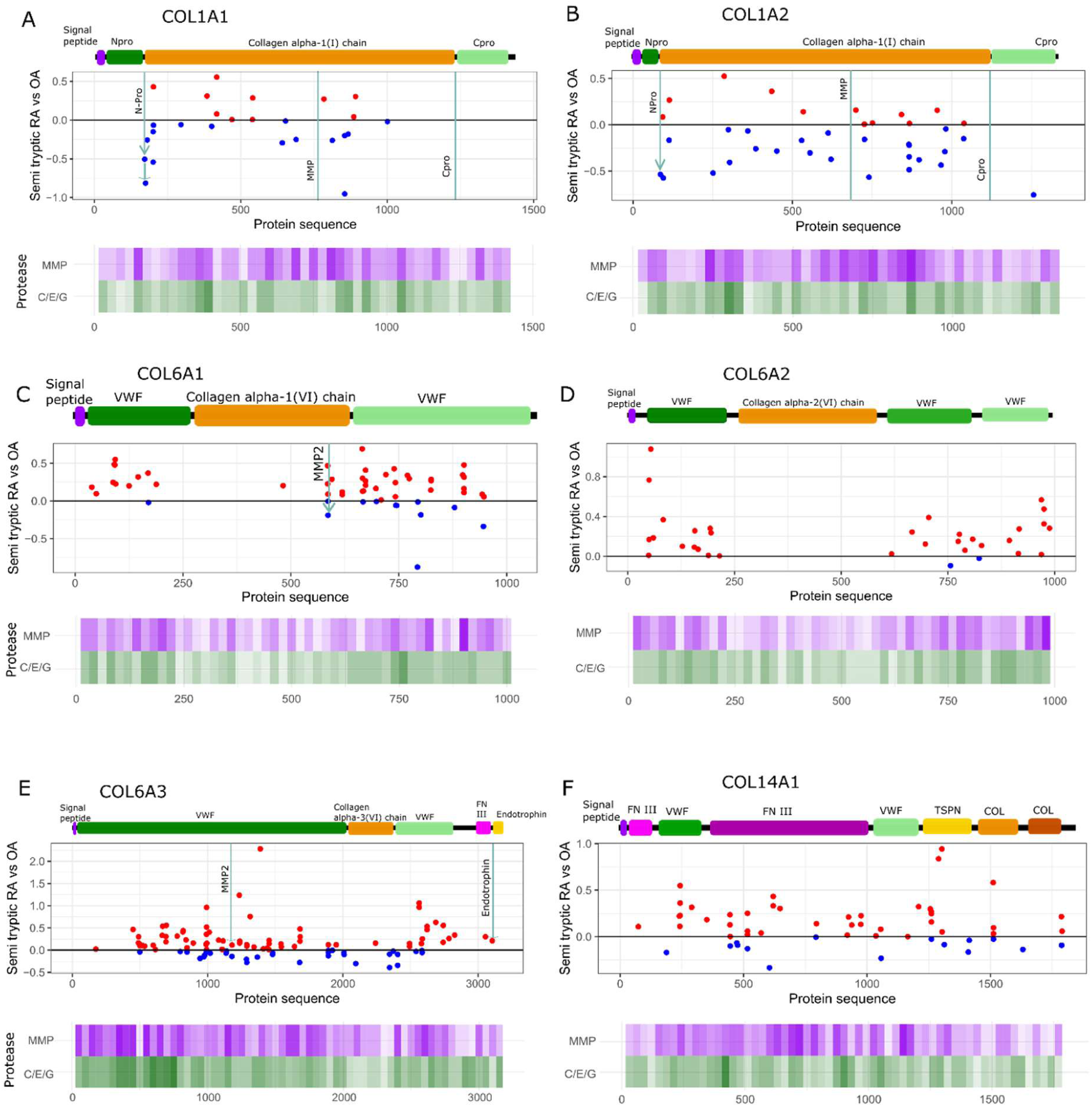
Mapping collagen cleavage sites in RA and OA. Fragment maps as shown in Figure 3C for (A) COL1A1 (B) COL1A2 (C) COL6A1 (D) COL6A2 (E) COL6A3 (F) COL14A1. VWF (Von Willebrand Factor domain, FN (Fibronectin type domain), TSPN (Thrombospondin-like, N terminal domain) and COL (Collagen like, triple-helical domain).

Protease susceptibility was mapped against cleavage sites to add confidence in this method of analysing protein degradation. Areas identified as the highest susceptibility to proteases match up to detected cleavage sites in every case for collagen-I, VI and XIV (the most degraded proteins identified in this study) (Figure 4). Degradation of these collagens also showed differences in RA and OA. Fragment abundances were normalised to their respective total protein abundance to focus on degradative ratios rather than changes in protein expression.

Collagen-I showed increased degradation in OA, whereas collagen-VI and XIV were more degraded in RA. These differences were also more pronounced in certain areas of the proteins. The first VWF region of collagen type VI alpha 1 (COL6A1) and the last VWF region of COL6A3 both appear more heavily favoured for RA degradation. Degradation of the first VWF domain of collagen-XIV is also heavily upregulated in RA.

Outside of collagens, domain specific cleavage patterns were detected for other matrix proteins. TNC and TNXB degradation seemed to favour fibronectin domains over epidermal growth factor (EGF)-like domains (Supplemental Figure 3I,J). Interestingly cleavage of the fibrinogen-like globule (FBG) domain was only detected in TNXB and was upregulated in OA compared to the rest of the protein which was more degraded in RA. MMP cleavage fragments were detected for core matrix proteins such as FN1, BGN and OGN (Supplemental Figure 3K-L). Further these fragments were all upregulated in RA. An additional protease cleavage site was clearly indicated in OGN with a cluster of detected fragments correlating with high protease susceptibility (Supplemental Figure 3L). In HSPG2, fragments matched to glycosylation sites, sugar additions would likely alter protease activity and so fragmentation differences between RA and OA may be reflective of differing glycosylation states (Supplementary Figure 3M). Fragments cleaved at N-linked glycosylation sites were upregulated in RA and those at O-linked sites were increased in OA.

### Semi-tryptic degradome analysis reveals RA endotype-specific fragments

Having demonstrated the disease specificity of the RA degradome we next sought to investigate the heterogeneity of RA pathology. RA can be stratified into distinct synovial endotypes based on immune cell infiltration, we focus here on myeloid and lymphoid pathotypes characterised by high proportions of macrophages and B/T cells respectively. Crucially these endotypes are predictive of differential treatment responses. Studies have found that patients with myeloid enriched disease respond better to tumor necrosis factor (TNF) blockade and those presenting with a lymphoid endotype responding better to anti-interleukin-6 receptor (IL-6R) therapy (*36*, *37*). Other investigations have shown a better response to rituximab, a B cell depletion therapy targeting CD20, in patients identified as lymphoid rather than myeloid RA (*38*, *39*). This difference in treatment response highlights the need for early and effective endotype diagnosis. To better understand whether the degradome was capable of distinguishing between disease endotypes we applied the DegrAID pipeline to a dataset of human synovial biopsies from RA patients displaying either a myeloid or lymphoid endotype (n=4) (*40*). 1790 matrisome fragments were detected, with over 600 of these resulting from both collagens (653) and glycoproteins (613). Overall counts showed similar trends to which previous datasets with high levels of degradation in collagens, fibronectin and fibrinogen (Figure 5A).

**Fig. 5.**
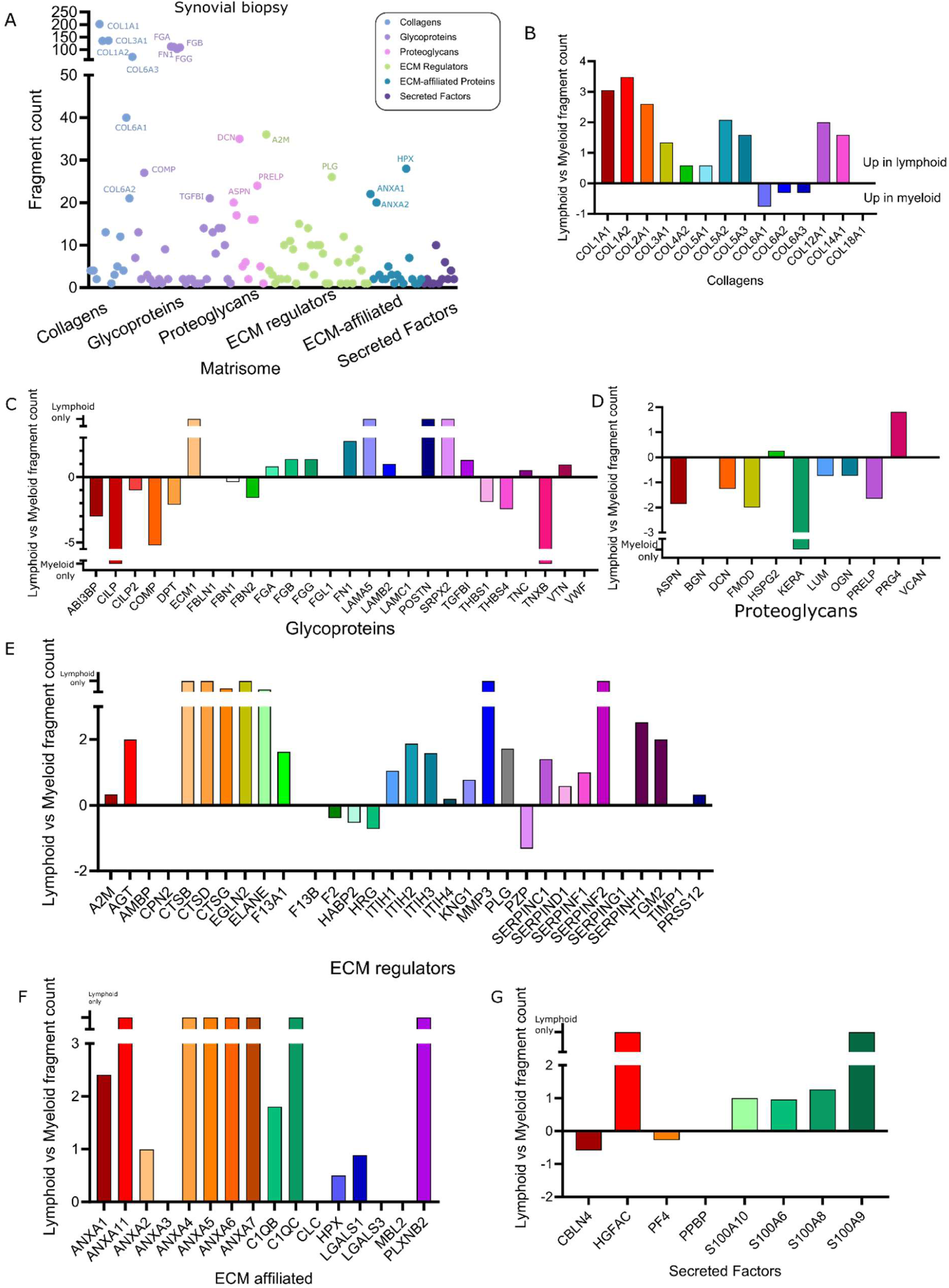
Comparison of RA endotype degradomes. (A) Total fragment counts for ECM proteins detected in synovial biopsies from RA patients coloured by their matrisome group. The 20 proteins with the most fragments were labelled. (B) Log fold change in fragment counts between myeloid and lymphoid endotypes for collagens. (C) As in (B) for glycoproteins. (D) As in (B) for proteoglycans. (E) As in (B) for ECM regulators. (F) As in (B) for ECM affiliated proteins. (G) As in (B) for secreted factors. Where breaks in bars are displayed fragments were only detected in one endotype.

To investigate endotype specific cleavages, fragment counts for individual proteins were analysed and compared (Figure 5B-G). Collagens largely displayed increased fragment counts in lymphoid samples and this was consistent across the group with the exception of collagen-VI which was more degraded in myeloid samples (Figure 5B). Cleavage maps further demonstrated the differences in collagen I and VI degradation between endotypes (Supplemental Figure 4A-E). Detected cleavage sites (orange for lymphoid and green for myeloid) were plotted along the sequence of these collagens. Collagen I showed a larger amount of fragmentation in lymphoid enriched disease, largely evenly distributed along their sequence. These again demonstrated very limited degradation in collagen-VI helical regions compared to collagen-I. Other matrix proteins were also differentially cleaved in lymphoid and myeloid endotypes (Figure 5B-F). Matrix associated proteins appeared generally more degraded in lymphoid phenotypes while proteoglycans were more degraded in myeloid enriched disease.

In order to explore the potential of this pipeline to discover endotype specific cleavage products, proteins which had dramatically more fragments for one of the endotypes were further investigated leading to the selection of eight proteins and 17 fragments to produce an endotype-specific fingerprint. Cartilage Intermediate Layer Protein (CILP), a cartilage scaffolding protein, and TNXB, both demonstrated myeloid specific degradation (Figure 6A,B). Cleavage sites were selected for the fingerprint if they were detected multiple times (+/-5 amino acids) adding to the confidence in detection. For both CILP and TNXB 3 cleavage sites reached this threshold.

**Fig. 6.**
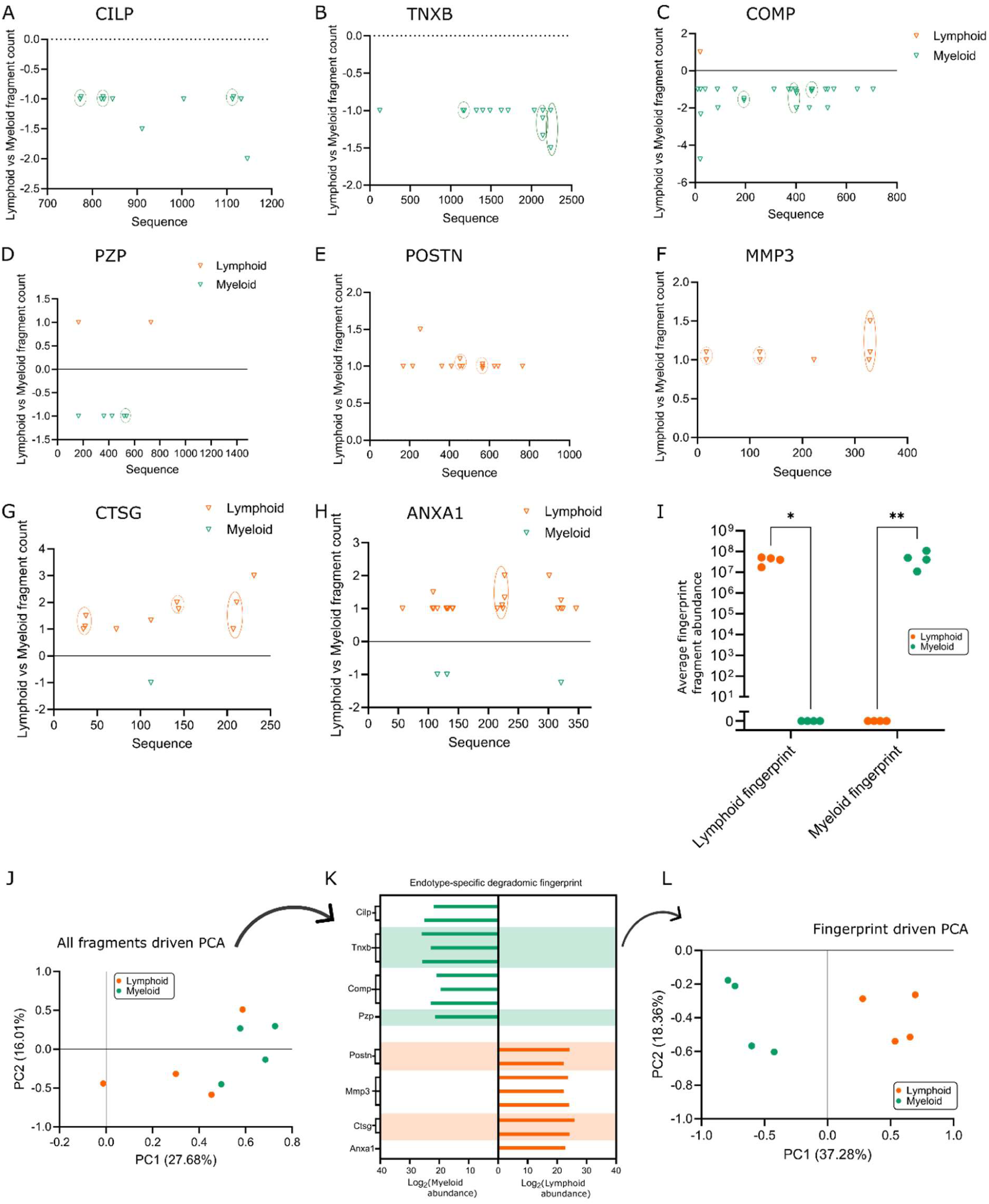
Endotype specific fragments. Endotype specific fragment map for (A) CILP, (B) TNXB, (C) COMP, (D) PZP, (E) POSTN, (F) MMP3, (G) CTSG, (H) ANXA1. Orange and green arrows indicate cleavage sites detected in lymphoid or myeloid endotype biopsies respectively. Height represents PSMs for identified fragments. Cleavage sites selected for fingerprint are circled. (I) Average abundance of lymphoid and myeloid fingerprints for lymphoid and myeloid biopsies. (J) Principal component analysis for the total degradome. (K) Endotype-specific degradomic fingerprint depicting average abundance of the different fragments included in the fingerprint. (L) Principal component analysis for the selected degradomic fingerprint.

Interestingly, no degradation was detected in the FBG domain, a region which is heavily conserved across the tenascin proteins, for TNXB in this dataset but FBG domain degradation was lymphoid specific in TNC (Supplementary Figure 5A). Degradation was substantially upregulated in the myeloid phenotype for both COMP (p=0.0004), a structural glycoprotein and pregnancy zone protein (PZP), a pan-protease inhibitor (p=0.07) (Figure 6C,D). These proteins provided 3 and 1 cleavage sites for the fingerprint respectively. Periostin (POSTN), a matricellular glycoprotein and stromelysin-1 (MMP3), a matrix metalloproteinase, showed lymphoid specific degradation generating 2 and 3 fragments for the fingerprints respectively (Figure 6E,F). Finally, we detected specific cleavages for 2 proteins with significantly increased degradation in the lymphoid endotype, cathepsin G (CTSG), a serine protease (p=0.002) and annexin A1 (ANXA1), a phospholipid binding, matrix associated protein (p<0.0001) (Figure 6G,H). Interestingly, this endotype specific cleavage of ANXA1 was not seen in Annexin A2.

Combining these identified cleavages allowed the generation of an endotype specific, degradomic fingerprint. The abundance of this fingerprint shows significant differences between myeloid and lymphoid patients (Figure 6I). The degradome as a whole was not able to separate endotypes using principal component analysis (Figure 6J). When the fingerprint described here is used, however, the patients clustered by endotype (Figure 6K,L). This demonstrated the potential of the degradome to generate disease and endotype specific fingerprints and moreover, this method of in-silico degradomic investigation.

We have shown that tissue datasets can be used to investigate and identify degradomic signatures. As the tissue is the where arthritic disease takes place, it makes sense to use this sample type to generate these fingerprints, biopsies are however invasive. We therefore sought to use a synovial fluid dataset to see whether these fragments were detectable in a more easily sampled sample type. A publicly available dataset containing synovial fluid samples from 18 RA patients was reanalysed using the DegrAID pipeline. These samples generated substantially different degradomic signatures whilst still detecting large numbers of semi-tryptic fragments (Supplementary Figure 5B). These fragments largely originated from ECM regulators and only low numbers of fragments resulting from collagens and proteoglycans were detected in comparison to tissue samples. This analysis identified 5 of the fragments identified in our degradomic signature (30%) demonstrating the potential to detect these peptides in more accessible fluids (Supplementary Figure 5C-H). This included 1 from COMP, POSTN and PZP and 2 from MMP3. 5 previously detected fragments from these proteins that were not selected for the fingerprint were also detected. In practice MS wouldn’t be used for this type of sampling due to the high cost and comparatively low throughput but this offers a proof of principal for fragment detection and the basis for development of more cost-effective ELISA based detection techniques.

## DISCUSSION

We have demonstrated how application of an in-silico analysis pipeline enables investigation of LC-MS/MS proteomics data generated without using specialist labelling methods to gain novel insights into tissue remodelling and degradation. Whilst this approach may not achieve the same depth as experimental degradomic approaches such as TAILS or HUNTER, it makes degradomic exploration hugely more accessible. Further, we show that it is both comparable to and correlates with data generated using fragment labelling and enrichment.

Comparison of tissues is challenging using LC-MS/MS proteomics, however interesting differences between osteoarthirtic synovial tissue and cartilage were identified. Whilst the majority of proteins showed similar trends towards either degradation or stability in both tissues, 18 didn’t. These included Collagen-II, COMP and ACAN all of which were identified as more actively degraded in cartilage. These are all key constituents of cartilage tissue that have been shown to decrease in osteoarthritic cartilage abundance as the disease progresses, suggesting an increase in their degradation as observed in this study (*31*, *41*, *42*). Collagen-VI, shown to be increasingly degraded in the synovium, has been shown to be highly expressed here, largely found in the synovial perivascular niche (*43*). Further to this, there is a distinction between fibrillar and non-fibrillar collagen degradation with the former showing cartilage-specific degradation and the latter being degraded in the synovium. Interestingly, key small leucine-rich proteoglycans (SLRPs), OGN, BGN, DCN, LUM and FMOD were all found to be degraded in both tissues which suggests a conserved pattern of SLRP turnover in the osteoarthritic joint. This fits with previous literature showing decreased SLRP abundance in OA and the detection of their cleavage products (*44*).

This approach also identified disease specific degradomes for RA and OA. The fact that these degradomes are clearly distinguishable opens the door for the potential use of degradomes in diagnosis more widely. Matrix fragments, termed matrikines, have previously been characterised both biologically and as potential biomarkers such as Endotrophin, a C-terminal fragment of Collagen alpha-3(VI). We detect Endotrophin fragments in both datasets and its cleavage appears increased in RA. Endotrophin has previously been shown to stimulate inflammation and fibrosis (*45*, *46*). Additionally, endotrophin serum levels have been shown to correlate with mortality in chronic kidney disease, heart failure and is predictive of treatment response in type 2 diabetes (*47–49*). Other collagen matrikines, restin and endostatin, C terminal fragments of Collagen XV and XVIII respectively and heavily conserved, both display anti-angiogenic properties (*50*). We detect cleavages within previously identified MMP sensitive regions that suggest release of these matrikines. Interestingly, treatment with Endostatin has been shown to reduce disease in collagen-induced arthritis mouse models, reducing synovial thickening and bone erosion (*51*). Matrikines also exist for non-collagenous matrix proteins such as Endorepellin, a C-terminal fragment of HSPG2, cleavage of which we also detect here, with increased abundance in RA. Endorepellin has been shown to be angiostatic, pro-autophagic and tumour-repressive (*52*). Additionally, it has demonstrated potential for use as a breast cancer biomarker (*53*). These matrikines demonstrate the important biological roles as well as potential use as biomarkers of other novel matrix fragments.

Clear differences in collagen-I and –VI degradation in RA and OA are identified in this study. Differential MMP landscapes between the diseases and variations in proteases which target these proteins could predict distinction between collagen degradomes. For example, MMP13, a protease important in OA, is known to cleave collagen-I and –II. MMP2 and 9, on the other hand, are both upregulated in active RA, in comparison to remission or OA, and are known to cleave collagen-VI (*43*).

Whilst collagen degradation fragments are not a new concept they have rarely been shown to distinguish between diseases or pathotypes as shown here, rather being used to track disease severity or treatment response. A collagen-I fragment termed C1M, for example, have been shown to increase in both RA and OA in comparison to healthy controls (*54*, *55*). An MMP driven collagen-VI fragment, C6M, has been shown to correlate with IL-6 receptor (Tocilizumab) treatment response in RA and whilst not disease-specific this does support the analysis in this study (*56*). Collagen VI degradation, through C6M detection, has been implicated in other diseases associated with high levels of tissue remodelling such as liver fibrosis, chronic obstructive pulmonary disease induced inflammation and ankylosing spondylitis (*57–59*).

Interestingly in RA, C-reactive protein (CRP), an established biomarker indicative of inflammation, wasn’t found to be differentially abundant between treatment responders and non-responders whereas C6M was, this suggests that using matrisomal, degradomic biomarkers may be more indicative of disease burden and long-term clinical response (*60*).

Collagen-VI degradation seen in this study also intriguingly appeared, in contrast to other collagens, to largely avoid triple helical regions instead accruing within VWF domains. These domains are known to be important for microfibril formation and mutations within these regions are associated with diseases such as muscular dystrophy and Bethlem Myopathy (*61*). Whilst specific proteases are known to be capable of cleaving the triple helical domain of collagen-I, allowing its unwinding before other gelatinases and cathepsins then further fragment the protein as we see evidence of, less is known about collagen-VI degradation. MMP9 mediated collagen-VI degradation, however, has been explored using mass spectrometry in combination with MMP9 knockdown. MMP9 sensitive regions of Collagen-VI were identified and all of these overlapped with VWF domains and areas with high levels of degradation identified in our study (*62*). As MMP9 abundance is increased in RA over OA this supports the substantial increase in degradation of collagen-VI particularly within these regions found in this study (*43*).

We also explored endotype specificity of the RA degradome. RA has a large degree of molecular and cellular heterogeneity; this can be characterised at a number of different levels, including histological description of two inflammatory phenotypes: myeloid and lymphoid. These are principally described by the diseased synovium being enriched for macrophages or B and T cells respectively and there are markers which can be used to distinguish subtypes such as sICAM1 for myeloid and CXCL13 for lymphoid (Dennis et al., 2014). These subtypes have also been shown to have different treatment responses to TNF, IL-6R and CD20 based therapies (*64*). We focus on the comparison of myeloid and lymphoid RA here to demonstrate the presence of endotype specific degradomes in synovial tissue and therefore the potential for degraded fragments to predict treatment response. Other endotypes have also been defined such as fibroid and pauci-immune and additional markers have been described (*65–67*). Further to this single cell technologies have been employed to characterise more in-depth subtypes of RA. This has led to the identification of cell-type abundance phenotypes (CTAPs) where patients were clustered based on enriched cell states which were found to correlate with cytokine and risk gene expression (*68*). Different CTAPs were also shown to respond to different treatments. Unfortunately, a lack of available proteomic data from RA patients with annotated CTAPs prevented analysis of these endotypes hence the focus here on myeloid and lymphoid RA.

We saw substantial differences in myeloid and lymphoid degradomes with interesting trends in collagens and proteoglycans. In the former, most collagens were more degraded in lymphoid endotypes with the exception of collagen-VI. Proteoglycans also appeared an exception to the majority of the matrisome, which displayed increased turnover in lymphoid, proteoglycans were instead largely degraded in the myeloid phenotype. We also identified 8 proteins which had endotype specific degradation. CILP, TNXB, COMP and PZP all had myeloid specific degradation fragments. CILP is a glycoprotein scaffolding protein which can be induced by transforming growth factor beta (TGFB). An MMP driven fragment has previously been identified to be increased in RA, and respond to TNF blockade, however this fragment does not match our identified cleavage sites (*69*).TNXB, a large glycoprotein with roles in tissue architecture and cell adhesion, has been shown to have differential methylation patterns in RA, alongside decreased gene expression (*69*). TXNB levels decrease in hemophilic arthropathy, another joint disease, with knockdown shown to cause ECM breakdown, this would predict that altered TXNB degradation would have a substantial impact on the tissue (*70*). We see extensive fragmentation of TXNB in the myeloid phenotype despite detecting none in lymphoid patients, all within Fibronectin type-III domains. We also identified heavy fragmentation of the glycoprotein COMP in myeloid patients. COMP protein is an established biomarker of RA with increased levels in both serum and synovial fluid which correlated with disease stage (*71*, *72*).

COMP fragments have also been shown to be increased in RA patient serum and synovial fluid (*73*, *74*). Further, MMPs associated with the myeloid endotype such as MMP19 have been shown to cleave COMP supporting the increased fragments detected in the myeloid phenotype here (*75*). The final myeloid fragment came from the protease PZP. This matrix regulator has been shown to increase in abundance in RA patient serum and decrease following Tocilizumab treatment, an IL-6R inhibitor (*76*, *77*).

The lymphoid specific degradomic fingerprint described in this study included fragments from POSTN, MMP3, CTSG and ANXA1. POSTN, a matrix glycoprotein, is largely degraded in the lymphoid subtype. Previously established to be increased in RA patients compared to OA in synovial tissue and fluid its levels also correlate with RA severity in lymphoid patients (*78*, *79*). Additionally, POSTN positive fibroblasts were increased in lymphoid patients which all supports the lymphoid specific degradation observed here. POSTN has also been shown to induce MMP3 expression in synovial cells, another protein for which we detect lymphoid specific fragments (*80*). As this would predict, especially alongside the fact it is largely produced by synovial fibroblasts and B cells, MMP protein has also been found to be elevated in both RA and specifically the lymphoid endotype (*81*, *82*). Moreover, MMP3 can predict radiographic disease outcome and treatment response (*82*, *83*). The cathepsin, CTSG, generated 3 lymphoid specific fragments and the protein has previously been shown to have increased abundance and activity in RA compared to OA (*84*, *85*). The final lymphoid fragment was of ANXA1, known to modulate the adaptive immune response it was been suggested as a possible RA therapeutic (*86*). ANXA1 levels have also been detected in peripheral CD4+ T cells in RA patients suggesting a lymphoid phenotype (*86*). Interestingly, ANXA1 is known to be cleaved by Proteinase-3 in the context of neutrophil activation and its fragments have been previously found to have roles in monocyte migration and are detected in cystic fibrosis patients (*87–89*). The proteins from which our degradomic fingerprint are generated all have supporting evidence for their involvement in RA, suggesting their turnover, as investigated here, would be important. The identified fragments however are novel and previously undescribed.

We detect many of these fragments using discovery proteomics in synovial fluid, which acts as a convincing proof of principle for the detection of these cleavage products in a more accessible, clinically relevant setting. ECM fragments have also previously been extensively shown to function as biomarkers (*22*, *90*, *91*). However, we nevertheless acknowledge that MS isn’t the best way to monitor these fragments and next steps would involve the development of specific screens so they are consistently detectable and at lower levels. The work here demonstrates the potential of degradomic investigation in tissue remodelling not only in identification of disease-or endotype-specific fragments but also to further our understanding of the biological implications of protein degradation. It is evident that matrix fragments are not produced randomly but instead are a result of specific and targeted degradation. Fragments can have functional roles as matrikines, they may feed into peptide generation for autoantibodies or lead to a loss or alteration to protein functioning or interaction through the destruction of specific protein domains.

Other limitations to this investigation largely revolve around reusing existing datasets. Running new degradomic analysis with fragment enrichment would increase both the numbers of degraded fragments identified and the confidence in their identification. This method however enables degradomic investigation without the need for cost and energy intensive rerunning of samples. If applied to the huge number of publicly accessible datasets and included in future analysis of proteomic experiments, its potential to further explore the degradome in a wide range of disease contexts is vast. Moreover, we have characterised RA synovial turnover, its degradative fingerprint and within that endotype specific degradomes, demonstrating that the RA degradome can bring to light an exciting new pool of clinically relevant targets.

## MATERIALS AND METHODS

Raw data was accessed from PRIDE, projects PXD28060, PXD021487, PXD027703, PXD044963 and PXD054336 and reanalysed using Fragpipe (v23). Data was searched against the UniProt/SwissProt human database (2024) with added decoys and known contaminants which contained a total of 40936 sequences (50% decoys) with a maximum false discovery rate (FDR) of 1%. Fixed modifications included in the search were carbamidomethyl addition to cysteine (+57.021Da) and, for relevant datasets, TMT6plex modifications at peptide N-termini and lysine residues (+229.163Da). Dynamic modifications of oxidation of methionine (+15.995Da), hydroxylation of proline (+15.995Da), deamination of asparagine/glutamine (+0.984) and phosphorylation of serine/threonine/tyrosine (+79.966) were also included. For TAILS analysis dimethyl modifications were included at peptide n-termini and lysine residue (+30.03Da). 2 missed cleavages were allowed. Both tryptic and semi-tryptic searches were run. Peptides were only included if they met the threshold 1% FDR and known contaminants were removed. Data was exported and onward analysis was conducted in Perseus (v2.0.11) and R (v4.3.2).

Semi-tryptic to tryptic ratios were calculated from average abundances of the relevant peptides for the protein of interest. Protein sequence mapping was done using semi-tryptic peptide abundance normalised to average tryptic peptide abundance. The Manchester Proteome was used to calculate protease susceptibility for two categories of protease, either MMPs 1,2,3,7,8,9,12,13,14 or Cathepsins b,d,g,k along with Elastase and Granzyme B. An MPC confidence of 0.5 was used in all cases with the frequency of cleavage plotted (*92*). Uniprot was used for domain information and Alphafold was used for protein structure predictions (*93*).

## List of Supplementary Materials

Fig S1 to S6

References and Notes

## Author contributions

Conceptualization: AH, KM Methodology and analysis: AH Funding acquisition: KM Writing: AH, KM

## Data and materials availability

All raw data was downloaded from publicly available datasets on the PRIDE repository, references to specific datasets can be found in the methods.

Any other data is available upon reasonable request.

## References

1. C. L. Bager, N. Willumsen, D. J. Leeming, V. Smith, M. A. Karsdal, D. Dornan, A. C. Bay-Jensen, Collagen degradation products measured in serum can separate ovarian and breast cancer patients from healthy controls: A preliminary study. Cancer Biomarkers 15, 783–788 (2015).

2. J. D. Hebert, S. A. Myers, A. Naba, G. Abbruzzese, J. M. Lamar, S. A. Carr, R. O. Hynes, Proteomic Profiling of the ECM of Xenograft Breast Cancer Metastases in Different Organs Reveals Distinct Metastatic Niches. Cancer Res. 80, 1475–1485 (2020).

3. M. A. Karsdal, V. B. Kraus, D. Shevell, A. C. Bay-Jensen, J. Schattenberg, R. Rambabu Surabattula, D. Schuppan, Profiling and targeting connective tissue remodeling in autoimmunity – A novel paradigm for diagnosing and treating chronic diseases. [Preprint] (2021). 10.1016/j.autrev.2020.102706.

4. X. Shao, C. D. Gomez, N. Kapoor, J. M. Considine, C. Grams, Y. (Tom) Gao, A. Naba, MatrisomeDB 2.0: 2023 updates to the ECM-protein knowledge database. Nucleic Acids Res. 51, D1519–D1530 (2023).

5. V. E. de Meijer, D. Y. Sverdlov, Y. Popov, H. D. Le, J. A. Meisel, V. Nosé, D. Schuppan, M. Puder, Broad-Spectrum Matrix Metalloproteinase Inhibition Curbs Inflammation and Liver Injury but Aggravates Experimental Liver Fibrosis in Mice. PLoS One 5, e11256 (2010).

6. R. Kalev-Altman, G. Becker, T. Levy, S. Penn, N. Y. Shpigel, E. Monsonego-Ornan, D. Sela-Donenfeld, Mmp2 Deficiency Leads to Defective Parturition and High Dystocia Rates in Mice. Int. J. Mol. Sci. 24, 16822 (2023).

7. R. Preston, R. Chrisp, M. Dudek, M. R. P. T. Morais, P. Tian, E. Williams, R. W. Naylor, B. Davenport, D. R. J. Pathiranage, E. Benson, D. G. Spiller, J. Bagnall, L. Zeef, C. Lawless, S. M. Baker, Q.-J. Meng, R. Lennon, The glomerular circadian clock temporally regulates basement membrane dynamics and the podocyte glucocorticoid response. Kidney Int. 107, 99–115 (2025).

8. J. Chang, R. Garva, A. Pickard, C.-Y. C. Yeung, V. Mallikarjun, J. Swift, D. F. Holmes, B. Calverley, Y. Lu, A. Adamson, H. Raymond-Hayling, O. Jensen, T. Shearer, Q. J. Meng, K. E. Kadler, Circadian control of the secretory pathway maintains collagen homeostasis. Nat. Cell Biol. 22, 74–86 (2020).

9. N. Yang, J. Williams, V. Pekovic-Vaughan, P. Wang, S. Olabi, J. McConnell, N. Gossan, A. Hughes, J. Cheung, C. H. Streuli, Q.-J. Meng, Cellular mechano-environment regulates the mammary circadian clock. Nat. Commun. 8, 14287 (2017).

10. T. Sasaki, N. Fukai, K. Mann, Structure, function and tissue forms of the C-terminal globular domain of collagen XVIII containing the angiogenesis inhibitor endostatin. EMBO J. 17, 4249–4256 (1998).

11. Y. Hamano, M. Zeisberg, H. Sugimoto, J. C. Lively, Y. Maeshima, C. Yang, R. O. Hynes, Z. Werb, A. Sudhakar, R. Kalluri, Physiological levels of tumstatin, a fragment of collagen IV α3 chain, are generated by MMP-9 proteolysis and suppress angiogenesis via αVβ3 integrin. Cancer Cell 3, 589–601 (2003).

12. A. Sudhakar, H. Sugimoto, C. Yang, J. Lively, M. Zeisberg, R. Kalluri, Human tumstatin and human endostatin exhibit distinct antiangiogenic activities mediated by αvβ3 and α5β1 integrins. Proceedings of the National Academy of Sciences 100, 4766–4771 (2003).

13. J. P. M. Blair, C. Bager, A. Platt, M. Karsdal, A.-C. Bay-Jensen, Identification of pathological RA endotypes using blood-based biomarkers reflecting tissue metabolism. A retrospective and explorative analysis of two phase III RA studies. PLoS One 14, e0219980 (2019).

14. K. Sun, J. Park, O. T. Gupta, W. L. Holland, P. Auerbach, N. Zhang, R. Goncalves Marangoni, S. M. Nicoloro, M. P. Czech, J. Varga, T. Ploug, Z. An, P. E. Scherer, Endotrophin triggers adipose tissue fibrosis and metabolic dysfunction. Nat. Commun. 5, 3485 (2014).

15. K. Henriksen, F. Genovese, A. Reese-Petersen, L. P. Audoly, K. Sun, M. A. Karsdal, P. E. Scherer, Endotrophin, a Key Marker and Driver for Fibroinflammatory Disease. Endocr. Rev. 45, 361–378 (2024).

16. T. Kantola, J. P. Väyrynen, K. Klintrup, J. Mäkelä, S. M. Karppinen, T. Pihlajaniemi, H. Autio-Harmainen, T. J. Karttunen, M. J. Mäkinen, A. Tuomisto, Serum endostatin levels are elevated in colorectal cancer and correlate with invasion and systemic inflammatory markers. Br. J. Cancer 111, 1605–1613 (2014).

17. M. S. O’Reilly, T. Boehm, Y. Shing, N. Fukai, G. Vasios, W. S. Lane, E. Flynn, J. R. Birkhead, B. R. Olsen, J. Folkman, Endostatin: An Endogenous Inhibitor of Angiogenesis and Tumor Growth. Cell 88, 277–285 (1997).

18. P. Garnero, S. Falkenløve Madsen, F. Eymard, J. Sellam, R. Chapurlat, A.-C. Bay-Jensen, Serum biochemical marker of synovial tissue turnover, C1M, predicts radiological progression in early rheumatoid arthritis. RMD Open 11, e005501 (2025).

19. S. Venkat, J. Rusbuldt, D. Richards, T. Freeman, C. Richmond, J. H. Mortensen, J. B. Seidelin, A. Poulsen, B. McRae, D. Ruane, Serum Collagen Biomarkers Are Reflective of Tissue Specific Fibroblasts Associated With Ulcerative Colitis Activity and Treatment Response to Ustekinumab. United European Gastroenterol. J. 13, 982–996 (2025).

20. B. Seeliger, J. Ruwisch, J. M. Bülow Sand, F. B. Simões, H. Jessen, E. Boerner, J. Fuge, K. Sewald, T. Welte, P. D. Wendel-Garcia, J. C. Schupp, D. J. Leeming, F. Bonella, A. Prasse, Differential Effects of Antifibrotic Treatment on Outcome Prediction via Serial Matrix Metalloproteinase-Degraded C-Reactive Protein Neoepitope Levels in Idiopathic Pulmonary Fibrosis. Chest, doi: 10.1016/j.chest.2025.08.047 (2025).

21. J. E. Collins, D. Robinson, N. Arden, A. C. Bay-Jensen, L. A. Deveza, M. Karsdal, C. Ladel, T. A. Perry, C. J. Swearingen, D. J. Hunter, V. B. Kraus, Biochemical biomarkers of knee osteoarthritis progression: Results from the FNIH biomarkers consortium progress OA study. Osteoarthr. Cartil. Open 7, 100677 (2025).

22. M.-A. Gkini, W. Liao, K. de Vlam, V. Chandran, S. H. Nielsen, Discovery and Clinical Validation of C1M and C4M as Soluble Biomarkers for Diagnosis, Prognosis, and Symptom Prediction in Psoriatic Disease and Other Inflammatory Arthropathies. J. Rheumatol. 52, 12–15 (2025).

23. M. Lindholm, L. E. Godskesen, T. Manon-Jensen, J. Kjeldsen, A. Krag, M. A. Karsdal, J. H. Mortensen, Endotrophin and C6Ma3, serological biomarkers of type VI collagen remodelling, reflect endoscopic and clinical disease activity in IBD. Sci. Rep. 11, 14713 (2021).

24. H. Port, B. Coppers, S. Tragl, E. Manger, L. M. Niemiec, S. Bayat, D. Simon, F. Fagni, G. Corte, A.-C. Bay-Jensen, K. Tascilar, A. J. Hueber, K. G. Schmidt, V. Schönau, M. Sticherling, S. Heinrich, S. Leyendecker, D. Bohr, G. Schett, A. Kleyer, S. Holm Nielsen, A.-M. Liphardt, Serum extracellular matrix biomarkers in rheumatoid arthritis, psoriatic arthritis and psoriasis and their association with hand function. Sci. Rep. 15, 13656 (2025).

25. S. Bhutada, L. Li, B. Willard, G. Muschler, N. Piuzzi, S. S. Apte, Forward and reverse degradomics defines the proteolytic landscape of human knee osteoarthritic cartilage and the role of the serine protease HtrA1. Osteoarthritis Cartilage 30, 1091–1102 (2022).

26. D. R. Martin, J. C. Witten, C. D. Tan, E. R. Rodriguez, E. H. Blackstone, G. B. Pettersson, D. E. Seifert, B. B. Willard, S. S. Apte, Proteomics identifies a convergent innate response to infective endocarditis and extensive proteolysis in vegetation components. JCI Insight 5 (2020).

27. S. Bhutada, A. Hoyle, N. S. Piuzzi, S. S. Apte, Degradomics defines proteolysis information flow from human knee osteoarthritis cartilage to matched synovial fluid and the contributions of secreted proteases ADAMTS5, MMP13 and CMA1 to articular cartilage breakdown. Osteoarthritis Cartilage 33, 116–127 (2025).

28. M. Ozols, A. Eckersley, C. I. Platt, C. Stewart-McGuinness, S. A. Hibbert, J. Revote, F. Li, C. E. M. Griffiths, R. E. B. Watson, J. Song, M. Bell, M. J. Sherratt, Predicting Proteolysis in Complex Proteomes Using Deep Learning. Int. J. Mol. Sci. 22, 3071 (2021).

29. J. Fleming, P. Magana, S. Nair, M. Tsenkov, D. Bertoni, I. Pidruchna, M. Q. Lima Afonso, A. Midlik, U. Paramval, A. Žídek, A. Laydon, O. Kovalevskiy, J. Pan, J. Cheng, Ž. Avsec, C. Bycroft, L. H. Wong, M. Last, M. Mirdita, M. Steinegger, P. Kohli, M. Váradi, S. Velankar, AlphaFold Protein Structure Database and 3D-Beacons: New Data and Capabilities. J. Mol. Biol. 437, 168967 (2025).

30. M. Cosenza-Contreras, A. Seredynska, D. Vogele, N. Pinter, E. Brombacher, R. F. Cueto, T. J. Dinh, P. Bernhard, M. Rogg, J. Liu, P. Willems, S. Stael, P. F. Huesgen, E. W. Kuehn, C. Kreutz, C. Schell, O. Schilling, TermineR: Extracting information on endogenous proteolytic processing from shotgun proteomics data. Proteomics 24 (2024).

31. M. R. Eveque-Mourroux, P. J. Emans, A. Boonen, B. S. R. Claes, F. G. Bouwman, R. M. A. Heeren, B. Cillero-Pastor, Heterogeneity of Lipid and Protein Cartilage Profiles Associated with Human Osteoarthritis with or without Type 2 Diabetes Mellitus. J. Proteome Res. 20, 2973–2982 (2021).

32. L. Alcaide-Ruggiero, R. Cugat, J. M. Domínguez, Proteoglycans in Articular Cartilage and Their Contribution to Chondral Injury and Repair Mechanisms. Int. J. Mol. Sci. 24, 10824 (2023).

33. R. de Groot, P. B. Folgado, K. Yamamoto, D. R. Martin, C. D. Koch, D. Debruin, S. Blagg, A. F. Minns, S. Bhutada, J. Ahnström, J. Larkin, A. Aspberg, P. Önnerfjord, S. S. Apte, S. Santamaria, Cleavage of Cartilage Oligomeric Matrix Protein (COMP) by ADAMTS4 generates a neoepitope associated with osteoarthritis and other forms of degenerative joint disease. Matrix Biology 135, 106–124 (2025).

34. X. Ren, M. Geng, K. Xu, C. Lu, Y. Cheng, L. Kong, Y. Cai, W. Hou, Y. Lu, Y. Aihaiti, P. Xu, Quantitative Proteomic Analysis of Synovial Tissue Reveals That Upregulated OLFM4 Aggravates Inflammation in Rheumatoid Arthritis. J. Proteome Res. 20, 4746–4757 (2021).

35. R. S. Pedersen, J. Thorlacius-Ussing, M. G. Raimondo, L. L. Langholm, G. Schett, A. Ramming, M. Karsdal, N. Willumsen, Fibroblast Activation Protein (FAP)-Mediated Cleavage of Type III Collagen Reveals Serum Biomarker Potential in Non-Small Cell Lung Cancer and Spondyloarthritis. Biomedicines 12, 545 (2024).

36. P. Juhl, A.-C. Bay-Jensen, M. Karsdal, A. S. Siebuhr, N. Franchimont, J. Chavez, Serum biomarkers of collagen turnover as potential diagnostic tools in diffuse systemic sclerosis: A cross-sectional study. PLoS One 13, e0207324 (2018).

37. N. Sparding, F. Genovese, M. A. Karsdal, N. M. Selby, Collagen type III formation but not degradation is associated with risk of kidney disease progression and mortality after acute kidney injury. Clin. Kidney J. 18 (2025).

38. G. Dennis, C. T. Holweg, S. K. Kummerfeld, D. F. Choy, A. F. Setiadi, J. A. Hackney, P. M. Haverty, H. Gilbert, W. Y. Lin, L. Diehl, S. Fischer, A. Song, D. Musselman, M. Klearman, C. Gabay, A. Kavanaugh, J. Endres, D. A. Fox, F. Martin, M. J. Townsend, Synovial phenotypes in rheumatoid arthritis correlate with response to biologic therapeutics. Arthritis Res. Ther. 16, R90 (2014).

39. J. Zhao, X. Ye, Z. Zhang, The predictive value of serum soluble ICAM-1 and CXCL13 in the therapeutic response to TNF inhibitor in rheumatoid arthritis patients who are refractory to csDMARDs. Clin. Rheumatol. 39, 2573–2581 (2020).

40. F. Rivellese, A. E. A. Surace, K. Goldmann, E. Sciacca, C. Çubuk, G. Giorli, C. R. John, A. Nerviani, L. Fossati-Jimack, G. Thorborn, M. Ahmed, E. Prediletto, S. E. Church, B. M. Hudson, S. E. Warren, P. M. McKeigue, F. Humby, M. Bombardieri, M. R. Barnes, M. J. Lewis, C. Pitzalis, F. Rivellese, G. Giorli, A. Nerviani, L. Fossati-Jimack, G. Thorborn, F. Humby, M. Bombardieri, M. J. Lewis, P. Durez, M. H. Buch, H. Rizvi, A. Mahto, C. Montecucco, B. Lauwerys, N. Ng, P. Ho, V. C. Romão, J. E. C. da Fonseca, P. Verschueren, S. Kelly, P. P. Sainaghi, N. Gendi, B. Dasgupta, A. Cauli, P. Reynolds, J. D. Cañete, J. Ramirez, R. Celis, R. Moots, P. C. Taylor, C. J. Edwards, J. Isaacs, P. Sasieni, E. Choy, C. Thompson, S. Bugatti, M. Bellan, M. Congia, C. Holroyd, A. Pratt, L. White, L. Warren, J. Peel, R. Hands, G. Hadfield, C. Pitzalis, Rituximab versus tocilizumab in rheumatoid arthritis: synovial biopsy-based biomarker analysis of the phase 4 R4RA randomized trial. Nat. Med. 28, 1256–1268 (2022).

41. S. Rosengren, N. Wei, K. C. Kalunian, A. Kavanaugh, D. L. Boyle, CXCL13: a novel biomarker of B-cell return following rituximab treatment and synovitis in patients with rheumatoid arthritis. Rheumatology 50, 603–610 (2011).

42. C. Çubuk, R. Lau, P. Cutillas, V. Rajeeve, C. R. John, A. E. A. Surace, R. Hands, L. Fossati-Jimack, M. J. Lewis, C. Pitzalis, Phosphoproteomic profiling of early rheumatoid arthritis synovium reveals active signalling pathways and differentiates inflammatory pathotypes. Arthritis Res. Ther. 26, 120 (2024).

43. S. Larsson, M. Englund, A. Struglics, L. S. Lohmander, Association between synovial fluid levels of aggrecan ARGS fragments and radiographic progression in knee osteoarthritis. Arthritis Res. Ther. 12, R230 (2010).

44. A. P. Hollander, T. F. Heathfield, C. Webber, Y. Iwata, R. Bourne, C. Rorabeck, A. R. Poole, Increased damage to type II collagen in osteoarthritic articular cartilage detected by a new immunoassay. Journal of Clinical Investigation 93, 1722–1732 (1994).

45. J.-B. Richard, A. Hoyle, M. Bower, S. Wu, L. Worthington, S. Davidson, Z. Varyova, C. Morrell, M. Pohin, B. Schonfeldova, Z. Y. Wong, L. MacDonald, M. Kurowska-Stolarska, S. G. Dakin, I. Udalova, C. A. Dendrou, A. Schwenzer, C. D. Buckley, K. S. Midwood, Synovial matrix turnover controls immune cell spatial patterning in inflammation resolution. Mol. Syst. Biol. 21, 1638–1665 (2025).

46. G.-X. Ni, Z. Li, Y.-Z. Zhou, The role of small leucine-rich proteoglycans in osteoarthritis pathogenesis. Osteoarthritis Cartilage 22, 896–903 (2014).

47. M. Kim, C. Lee, D. Y. Seo, H. Lee, J. D. Horton, J. Park, P. E. Scherer, The impact of endotrophin on the progression of chronic liver disease. Exp. Mol. Med. 52, 1766–1776 (2020).

48. K. M. Mak, C. Y. M. Png, “Type VI Collagen: Biological Functions and Its Neo-epitope as Hepatic Fibrosis Biomarker” (2015), pp. 1–27.

49. A. Fenton, M. D. Jesky, C. J. Ferro, J. Sørensen, M. A. Karsdal, P. Cockwell, F. Genovese, Serum endotrophin, a type VI collagen cleavage product, is associated with increased mortality in chronic kidney disease. PLoS One 12, e0175200 (2017).

50. J. A. Chirinos, L. Zhao, A. L. Reese-Petersen, J. B. Cohen, F. Genovese, A. M. Richards, R. N. Doughty, J. Díez, A. González, R. Querejeta, P. Zamani, J. Nuñez, Z. Wang, C. Ebert, K. Kammerhoff, J. Maranville, M. Basso, C. Qian, D. G. K. Rasmussen, P. H. Schafer, D. Seiffert, M. A. Karsdal, D. A. Gordon, F. Ramirez-Valle, T. P. Cappola, Endotrophin, a Collagen VI Formation–Derived Peptide, in Heart Failure. NEJM Evidence 1 (2022).

51. M. A. Karsdal, K. Henriksen, F. Genovese, D. J. Leeming, M. J. Nielsen, B. J. Riis, C. Christiansen, I. Byrjalsen, D. Schuppan, Serum endotrophin identifies optimal responders to PPARγ agonists in type 2 diabetes. Diabetologia 60, 50–59 (2017).

52. S. Bretaud, E. Guillon, S.-M. Karppinen, T. Pihlajaniemi, F. Ruggiero, Collagen XV, a multifaceted multiplexin present across tissues and species. Matrix Biol. Plus 6–7, 100023 (2020).

53. D. Kurosaka, K. Yoshida, J. Yasuda, T. Yokoyama, I. Kingetsu, N. Yamaguchi, K. Joh, M. Matsushima, S. Saito, A. Yamada, Inhibition of arthritis by systemic administration of endostatin in passive murine collagen induced arthritis. Ann. Rheum. Dis. 62, 677–679 (2003).

54. G. Bix, R. Castello, M. Burrows, J. J. Zoeller, M. Weech, R. A. Iozzo, C. Cardi, M. L. Thakur, C. A. Barker, K. Camphausen, R. V. Iozzo, Endorepellin In Vivo: Targeting the Tumor Vasculature and Retarding Cancer Growth and Metabolism. JNCI: Journal of the National Cancer Institute 98, 1634–1646 (2006).

55. J. W. Chang, U. Kang, D. H. Kim, J. K. Yi, J. W. Lee, D. Noh, C. Lee, M. Yu, Identification of circulating endorepellin LG3 fragment: Potential use as a serological biomarker for breast cancer. Proteomics Clin. Appl. 2, 23–32 (2008).

56. K. I. Maijer, N. S. Gudmann, M. A. Karsdal, D. M. Gerlag, P. P. Tak, A. C. Bay-Jensen, Neo-Epitopes—Fragments of Cartilage and Connective Tissue Degradation in Early Rheumatoid Arthritis and Unclassified Arthritis. PLoS One 11, e0149329 (2016).

57. A. C. Bay-Jensen, A. Mobasheri, C. S. Thudium, V. B. Kraus, M. A. Karsdal, Blood and urine biomarkers in osteoarthritis – an update on cartilage associated type II collagen and aggrecan markers. Curr. Opin. Rheumatol. 34, 54–60 (2022).

58. C. S. Thudium, P. Frederiksen, M. A. Karsdal, A.-C. Bay-Jensen, Changes in type VI collagen degradation reflect clinical response to treatment in rheumatoid arthritis patients treated with tocilizumab. Arthritis Res. Ther. 26, 3 (2024).

59. A. C. Bay-Jensen, D. J. Leeming, A. Kleyer, S. S. Veidal, G. Schett, M. A. Karsdal, Ankylosing spondylitis is characterized by an increased turnover of several different metalloproteinase-derived collagen species: a cross-sectional study. Rheumatol. Int. 32, 3565–3572 (2012).

60. N. Barascuk, S. S. Veidal, L. Larsen, D. V. Larsen, M. R. Larsen, J. Wang, Q. Zheng, R. Xing, Y. Cao, L. M. Rasmussen, M. A. Karsdal, A novel assay for extracellular matrix remodeling associated with liver fibrosis: An enzyme-linked immunosorbent assay (ELISA) for a MMP-9 proteolytically revealed neo-epitope of type III collagen. Clin. Biochem. 43, 899–904 (2010).

61. A. R. Bihlet, M. A. Karsdal, J. M. B. Sand, D. J. Leeming, M. Roberts, W. White, R. Bowler, Biomarkers of extracellular matrix turnover are associated with emphysema and eosinophilic-bronchitis in COPD. Respir. Res. 18, 22 (2017).

62. A. C. Bay-Jensen, A. Platt, I. Byrjalsen, P. Vergnoud, C. Christiansen, M. A. Karsdal, Effect of tocilizumab combined with methotrexate on circulating biomarkers of synovium, cartilage, and bone in the LITHE study. Semin. Arthritis Rheum. 43, 470–478 (2014).

63. H. Solomon-Degefa, J. M. Gebauer, C. M. Jeffries, C. D. Freiburg, P. Meckelburg, L. E. Bird, U. Baumann, D. I. Svergun, R. J. Owens, J. M. Werner, E. Behrmann, M. Paulsson, R. Wagener, Structure of a collagen VI α3 chain VWA domain array: adaptability and functional implications of myopathy causing mutations. Journal of Biological Chemistry 295, 12755–12771 (2020).

64. D. Xu, N. Suenaga, M. J. Edelmann, R. Fridman, R. J. Muschel, B. M. Kessler, Novel MMP-9 substrates in cancer cells revealed by a label-free quantitative proteomics approach. Mol. Cell. Proteomics 7, 2215–28 (2008).

65. G. Dennis, C. T. Holweg, S. K. Kummerfeld, D. F. Choy, A. F. Setiadi, J. A. Hackney, P. M. Haverty, H. Gilbert, W. Y. Lin, L. Diehl, S. Fischer, A. Song, D. Musselman, M. Klearman, C. Gabay, A. Kavanaugh, J. Endres, D. A. Fox, F. Martin, M. J. Townsend, Synovial phenotypes in rheumatoid arthritis correlate with response to biologic therapeutics. Arthritis Res. Ther. 16, R90 (2014).

66. J. Zhao, S. Guo, S. J. Schrodi, D. He, Molecular and Cellular Heterogeneity in Rheumatoid Arthritis: Mechanisms and Clinical Implications. Front. Immunol. 12, 790122 (2021).

67. G. Dennis, C. T. Holweg, S. K. Kummerfeld, D. F. Choy, A. F. Setiadi, J. A. Hackney, P. M. Haverty, H. Gilbert, W. Y. Lin, L. Diehl, S. Fischer, A. Song, D. Musselman, M. Klearman, C. Gabay, A. Kavanaugh, J. Endres, D. A. Fox, F. Martin, M. J. Townsend, Synovial phenotypes in rheumatoid arthritis correlate with response to biologic therapeutics. Arthritis Res. Ther. 16, R90 (2014).

68. G. Lliso-Ribera, F. Humby, M. Lewis, A. Nerviani, D. Mauro, F. Rivellese, S. Kelly, R. Hands, F. Bene, N. Ramamoorthi, J. A. Hackney, A. Cauli, E. H. Choy, A. Filer, P. C. Taylor, I. McInnes, M. J. Townsend, C. Pitzalis, Synovial tissue signatures enhance clinical classification and prognostic/treatment response algorithms in early inflammatory arthritis and predict requirement for subsequent biological therapy: results from the pathobiology of early arthritis cohort (PEAC). Ann. Rheum. Dis. 78, 1642–1652 (2019).

69. H. Johnsson, A. Najm, Synovial biopsies in clinical practice and research: current developments and perspectives. Clin. Rheumatol. 40, 2593–2600 (2021).

70. F. Zhang, A. H. Jonsson, A. Nathan, N. Millard, M. Curtis, Q. Xiao, M. Gutierrez-Arcelus, W. Apruzzese, G. F. M. Watts, D. Weisenfeld, S. Nayar, J. Rangel-Moreno, N. Meednu, K. E. Marks, I. Mantel, J. B. Kang, L. Rumker, J. Mears, K. Slowikowski, K. Weinand, D. E. Orange, L. Geraldino-Pardilla, K. D. Deane, D. Tabechian, A. Ceponis, G. S. Firestein, M. Maybury, I. Sahbudin, A. Ben-Artzi, A. M. Mandelin, A. Nerviani, M. J. Lewis, F. Rivellese, C. Pitzalis, L. B. Hughes, D. Horowitz, E. DiCarlo, E. M. Gravallese, B. F. Boyce, J. Albrecht, J. L. Barnas, J. M. Bathon, D. L. Boyle, S. L. Bridges, D. Campbell, H. L. Carr, A. Chicoine, A. Cordle, P. Dunn, L. Forbess, P. K. Gregersen, J. M. Guthridge, L. B. Ivashkiv, K. Ishigaki, J. A. James, G. Keras, I. Korsunsky, A. Lakhanpal, J. A. Lederer, Z. J. Li, Y. Li, A. McDavid, M. J. McGeachy, K. Raza, Y. Reshef, C. Ritchlin, W. H. Robinson, S. Sakaue, J. A. Seifert, A. Singaraju, M. H. Smith, D. Scheel-Toellner, P. J. Utz, M. H. Weisman, A. Wyse, Z. Zhu, L. W. Moreland, S. M. Goodman, H. Perlman, V. M. Holers, K. P. Liao, A. Filer, V. P. Bykerk, K. Wei, D. A. Rao, L. T. Donlin, J. H. Anolik, M. B. Brenner, S. Raychaudhuri, Deconstruction of rheumatoid arthritis synovium defines inflammatory subtypes. Nature 623, 616–624 (2023).

71. H. Port, C. M. Hausgaard, Y. He, W. P. Maksymowych, S. Wichuk, D. Sinkeviciute, A.-C. Bay-Jensen, S. Holm Nielsen, A novel biomarker of MMP-cleaved cartilage intermediate layer protein-1 is elevated in patients with rheumatoid arthritis, ankylosing spondylitis and osteoarthritis. Sci. Rep. 13, 21717 (2023).

72. J. Chen, Q. Zeng, X. Wang, R. Xu, W. Wang, Y. Huang, Q. Sun, W. Yuan, P. Wang, D. Chen, P. Tong, H. Jin, Aberrant methylation and expression of TNXB promote chondrocyte apoptosis and extracullar matrix degradation in hemophilic arthropathy via AKT signaling. Elife 13 (2024).

73. T. Saxne, D. Heinegård, Cartilage Oligomeric Matrix Protein: A novel marker of cartilage turnover detectable in synovial fluid and blood. Rheumatology 31, 583–591 (1992).

74. F. Liu, X. Wang, X. Zhang, C. Ren, J. Xin, Role of Serum cartilage oligomeric matrix protein (COMP) in the diagnosis of rheumatoid arthritis (RA): A case–control study. Journal of International Medical Research 44, 940–949 (2016).

75. E. Åhrman, P. Lorenzo, K. Holmgren, A. J. Grodzinsky, L. E. Dahlberg, T. Saxne, D. Heinegård, P. Önnerfjord, Novel Cartilage Oligomeric Matrix Protein (COMP) Neoepitopes Identified in Synovial Fluids from Patients with Joint Diseases Using Affinity Chromatography and Mass Spectrometry. Journal of Biological Chemistry 289, 20908–20916 (2014).

76. M. Neidhart, N. Hauser, M. Paulsson, P. E. DiCesare, B. A. Michel, H. J. Hauselmann, Small fragments of cartilage oligomeric matrix protein in synovial fluid and serum as markers for cartilage degradation. Rheumatology 36, 1151–1160 (1997).

77. J. O. Stracke, A. J. Fosang, K. Last, F. A. Mercuri, A. M. Pendás, E. Llano, R. Perris, P. E. Di Cesare, G. Murphy, V. Knäuper, Matrix metalloproteinases 19 and 20 cleave aggrecan and cartilage oligomeric matrix protein (COMP). FEBS Lett. 478, 52–56 (2000).

78. C. H. Horne, A. W. Thomson, C. B. Hunter, A. M. Tunstall, C. M. Towler, M. E. Billingham, Pregnancy-associated alpha 2-glycoprotein (alpha 2-PAG) and various acute phase reactants in rheumatoid arthritis and osteoarthritis. Biomedicine 30, 90–4 (1979).

79. M. Yanagida, M. Kawasaki, M. Fujishiro, M. Miura, K. Ikeda, K. Nozawa, H. Kaneko, S. Morimoto, Y. Takasaki, H. Ogawa, K. Takamori, I. Sekigawa, Serum Proteome Analysis in Patients with Rheumatoid Arthritis Receiving Therapy with Tocilizumab: An Anti-Interleukin-6 Receptor Antibody. Biomed Res. Int. 2013, 1–9 (2013).

80. R. Micheroli, M. Elhai, S. Edalat, M. Frank-Bertoncelj, K. Bürki, A. Ciurea, L. MacDonald, M. Kurowska-Stolarska, M. J. Lewis, K. Goldmann, C. Cubuk, T. Kuret, O. Distler, C. Pitzalis, C. Ospelt, Role of synovial fibroblast subsets across synovial pathotypes in rheumatoid arthritis: a deconvolution analysis. RMD Open 8, e001949 (2022).

81. N. Athanasou, A. Mantoku, A. Kudo, Y. Inagaki, D. Mahoney, T. Kashima, Periostin expression in inflammatory and non-inflammatory osteoarthritis and rheumatoid arthritis. Osteoarthritis Cartilage 24, S69 (2016).

82. Y. Tajika, T. Moue, S. Ishikawa, K. Asano, T. Okumo, H. Takagi, T. Hisamitsu, Influence of Periostin on Synoviocytes in Knee Osteoarthritis. In Vivo 31, 69–77 (2017).

83. F. Humby, M. Lewis, N. Ramamoorthi, J. A. Hackney, M. R. Barnes, M. Bombardieri, A. F. Setiadi, S. Kelly, F. Bene, M. DiCicco, S. Riahi, V. Rocher, N. Ng, I. Lazarou, R. Hands, D. van der Heijde, R. B. M. Landewé, A. van der Helm-van Mil, A. Cauli, I. McInnes, C. D. Buckley, E. H. Choy, P. C. Taylor, M. J. Townsend, C. Pitzalis, Synovial cellular and molecular signatures stratify clinical response to csDMARD therapy and predict radiographic progression in early rheumatoid arthritis patients. Ann. Rheum. Dis. 78, 761–772 (2019).

84. A. Lerner, S. Neidhöfer, S. Reuter, T. Matthias, MMP3 is a reliable marker for disease activity, radiological monitoring, disease outcome predictability, and therapeutic response in rheumatoid arthritis. Best Pract. Res. Clin. Rheumatol. 32, 550–562 (2018).

85. M. Houseman, C. Potter, N. Marshall, R. Lakey, T. Cawston, I. Griffiths, S. Young-Min, J. D. Isaacs, Baseline serum MMP-3 levels in patients with Rheumatoid Arthritis are still independently predictive of radiographic progression in a longitudinal observational cohort at 8 years follow up. Arthritis Res. Ther. 14, R30 (2012).

86. D. Trzybulska, A. Olewicz-Gawlik, K. Graniczna, K. Kisiel, M. Moskal, D. Cieślak, P. Hrycaj, Quantitative analysis of elastase and cathepsin G mRNA levels in peripheral blood CD14+ cells from patients with rheumatoid arthritis. Cell. Immunol. 292, 40–44 (2014).

87. J. Miyata, K. Tani, K. Sato, S. Otsuka, T. Urata, B. Lkhagvaa, C. Furukawa, N. Sano, S. Sone, Cathepsin G: the significance in rheumatoid arthritis as a monocyte chemoattractant. Rheumatol. Int. 27, 375–382 (2007).

88. F. D’Acquisto, N. Paschalidis, K. Raza, C. D. Buckley, R. J. Flower, M. Perretti, Glucocorticoid treatment inhibits annexin-1 expression in rheumatoid arthritis CD4+ T cells. Rheumatology 47, 636–639 (2008).

89. L. Vong, F. D’Acquisto, M. Pederzoli-Ribeil, L. Lavagno, R. J. Flower, V. Witko-Sarsat, M. Perretti, Annexin 1 Cleavage in Activated Neutrophils. Journal of Biological Chemistry 282, 29998–30004 (2007).

90. K. E. Blume, S. Soeroes, H. Keppeler, S. Stevanovic, D. Kretschmer, M. Rautenberg, S. Wesselborg, K. Lauber, Cleavage of Annexin A1 by ADAM10 during Secondary Necrosis Generates a Monocytic “Find-Me” Signal. The Journal of Immunology 188, 135–145 (2012).

91. M. Perretti, R. J. Flower, Annexin 1 and the biology of the neutrophil. J. Leukoc. Biol. 76, 25–29 (2004).

92. J. E. Collins, D. Robinson, N. Arden, A. C. Bay-Jensen, L. A. Deveza, M. Karsdal, C. Ladel, T. A. Perry, C. J. Swearingen, D. J. Hunter, V. B. Kraus, Biochemical biomarkers of knee osteoarthritis progression: Results from the FNIH biomarkers consortium progress OA study. Osteoarthr. Cartil. Open 7, 100677 (2025).

93. J. M. B. Sand, P. Frederiksen, A. E. John, F. B. Simões, N. Hoyer, T. S. Prior, A. Avdic-Belltheus, P. L. Molyneaux, I. D. Stewart, H. P. Fainberg, S. R. Johnson, M. A. Karsdal, D. J. Leeming, E. Bendstrup, S. B. Shaker, T. M. Maher, G. Jenkins, Basement membrane repair response biomarker PRO-C4 predicts progression in idiopathic pulmonary fibrosis: analysis of the PFBIO and PROFILE cohorts. Thorax 80, 935–944 (2025).

