## Supplemental Figures for "In silico degradomics reveals disease- and endotype-specific alterations in the joint tissue landscape"

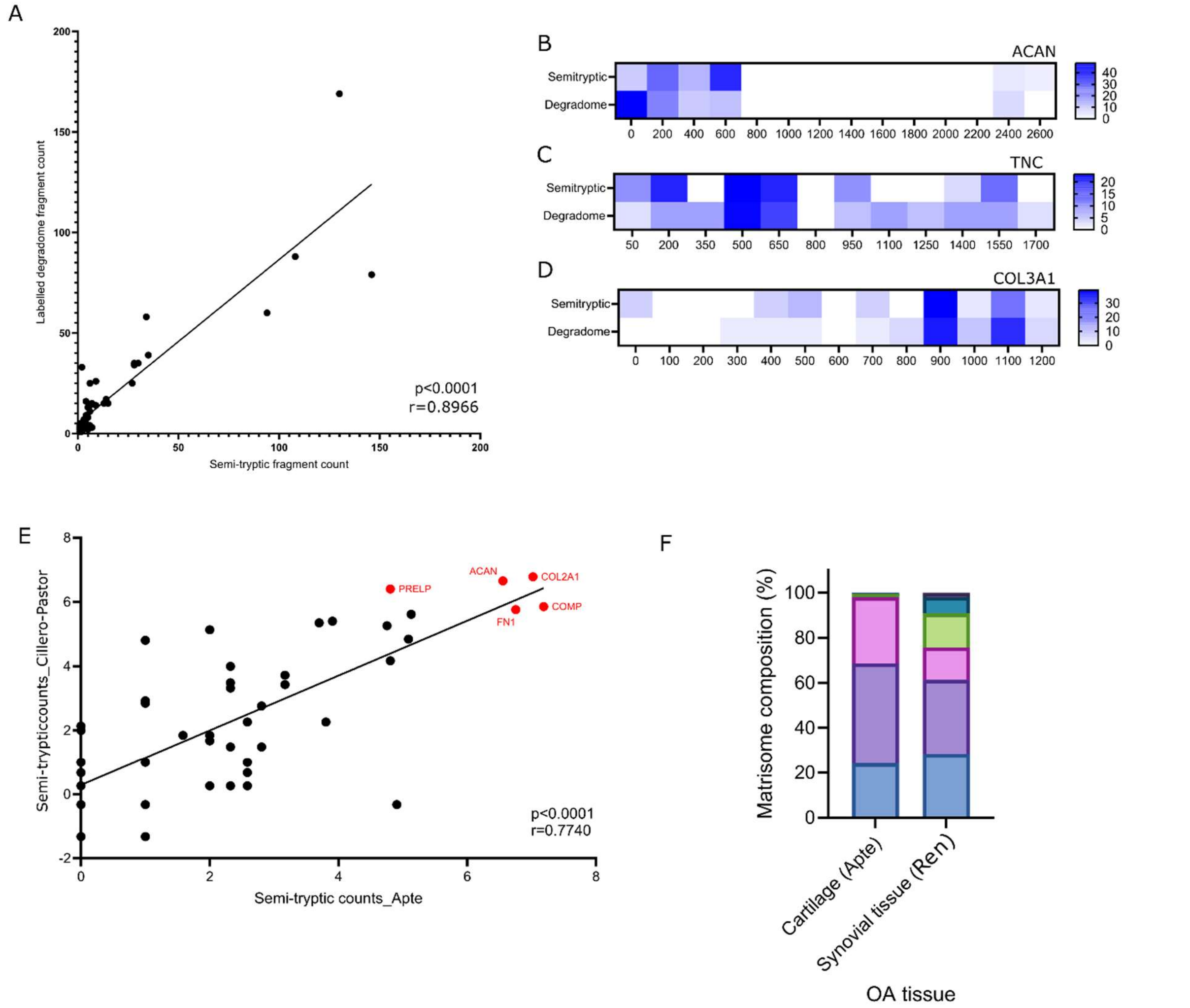


**Supplementary Fig. 1.**
(A) Correlation of fragment counts resulting from either TAILS or semi-tryptic degradomic analysis. Significant association was determined by Pearson correlation ($r=0.8966$, $p<0.0001$). (B) Fragment map for ACAN, coloured by numbers of cleavage sites detected per 200 amino acids by semi-tryptic or TAILS analysis. (C) As in (B) for TNC, every 150 amino acids. (D) As in (B) for COL3A1 every 100 amino acids. (E) Correlation of fragment counts between Apte and Cillero-Pastor labs for OA cartilage tissue. Significant association was determined by Pearson correlation ($r=0.7740$, $p<0.0001$). (F) Relative matrisomal abundance of cartilage and synovial tissue OA degradomes. Matrisome groups determined by MatrisomeDB.

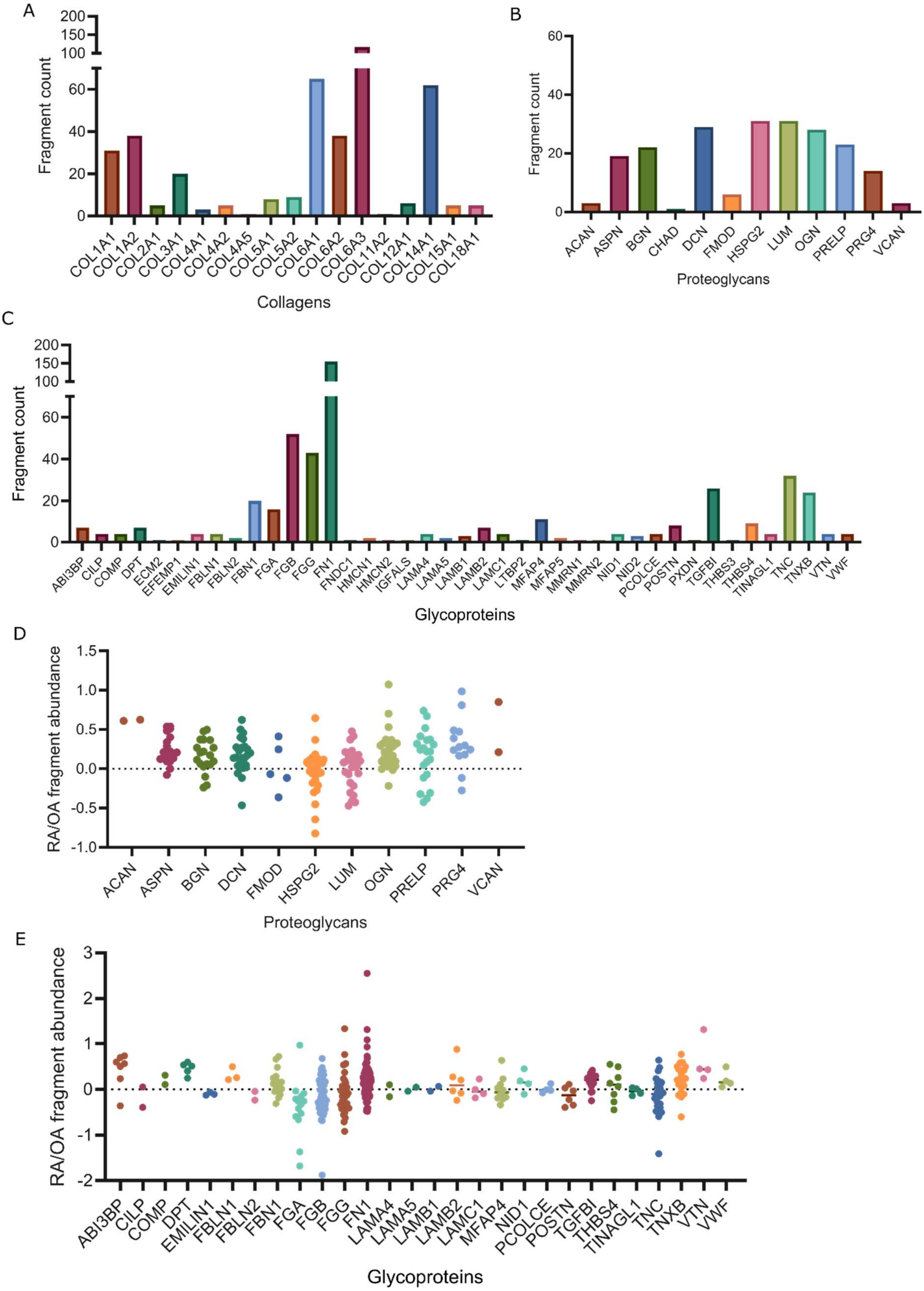
A
Fragment count
200
150
100
60
40
20
0
COL1A1
COL1A2
COL2A1
COL3A1
COL4A1
COL4A2
COL4A5
COL5A1
COL5A2
COL6A1
COL6A2
COL6A3
COL11A2
COL12A1
COL14A1
COL15A1
COL18A1
Collagens
B
Fragment count
60
40
20
0
ACAN
ASPN
BGN
CHAD
DCN
FMOD
HSPG2
LUM
OGN
PRELP
PRG4
VCAN
Proteoglycans
C
Fragment count
200
150
100
60
40
20
0
ABI3BP
CILP
COMP
DPT
ECM2
EFEMP1
EMILIN1
FBLN1
FBLN2
FBN1
FGA
FGB
FGG
FN1
FNDC1
HMCN1
HMCN2
IGFALS
LAMA4
LAMA5
LAMB1
LAMB2
LAMC1
LTBP2
MFAP4
MFAP5
MMRN1
MMRN2
NID1
NID2
PCOLCE
POSTN
PXDN
TGFBI
THBS3
THBS4
TINAGL1
TNC
TNXB
VTN
VWF
Glycoproteins
D
RA/OA fragment abundance
1.5
1.0
0.5
0.0
-0.5
-1.0
ACAN
ASPN
BGN
DCN
FMOD
HSPG2
LUM
OGN
PRELP
PRG4
VCAN
Proteoglycans
E
RA/OA fragment abundance
3
2
1
0
-1
-2
ABI3BP
CILP
COMP
DPT
EMILIN1
FBLN1
FBLN2
FBN1
FGA
FGB
FGG
FN1
LAMA4
LAMA5
LAMB1
LAMB2
LAMC1
MFAP4
NID1
PCOLCE
POSTN
TGFBI
THBS4
TINAGL1
TNC
TNXB
VTN
VWF
Glycoproteins

**Supplementary Fig. 2.**

Semi-tryptic fragment counts in RA/OA dataset for (A) Collagens, (B) Proteoglycans, (C) Glycoproteins. (D) Proteoglycan fragment abundance (Log2) in RA compared to OA patients. Each dot relates to a different semi-tryptic peptide detected. (E) As in (D) for glycoproteins.

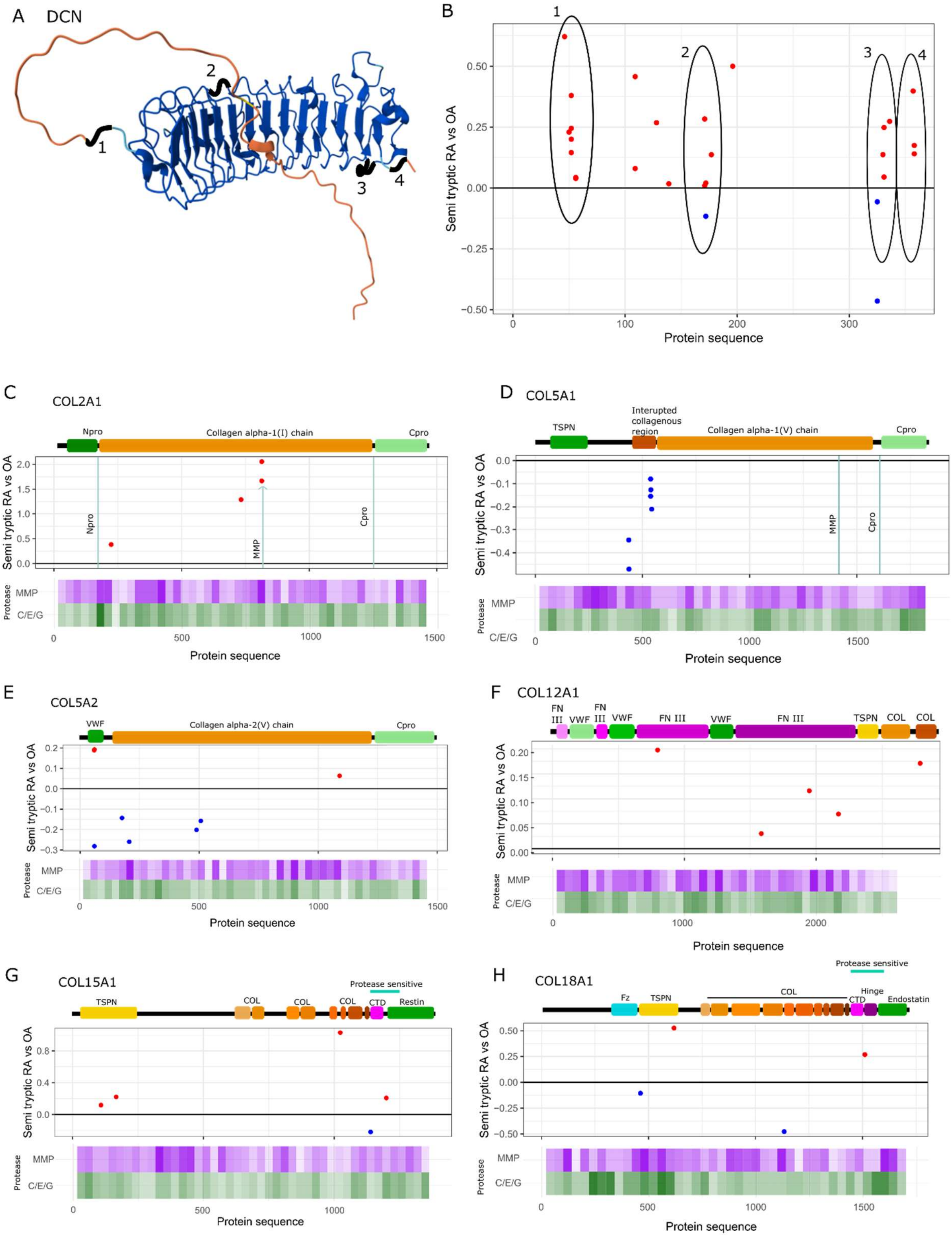

A DCN
1
2
3
4
B
Semi tryptic RA vs OA
Protein sequence
C COL2A1
Npro
Collagen alpha-1(I) chain
Cpro
MMP
C/E/G
Protease
D COL5A1
TSPN
Interupted collagenous region
Collagen alpha-1(V) chain
E COL5A2
VWF
Collagen alpha-2(V) chain
F COL12A1
FN III
VWF
TSPN
COL
G COL15A1
Protease sensitive
CTD
Restin
H COL18A1
Fz
Hinge
Endostatin

**Supplementary Fig. 3.**

(A) Alphafold predicted structure for DCN. Black highlighted sections correlate to cleavage sites detected in (B). (B) Cleavage map for DCN plotted as in Figure 3C. (C) Cleavage map as in 3C for COL2A1, (D) for COL5A1, (E) for COL5A2, (F) for COL12A1, (G) for COL15A1, (H) for COL18A1.

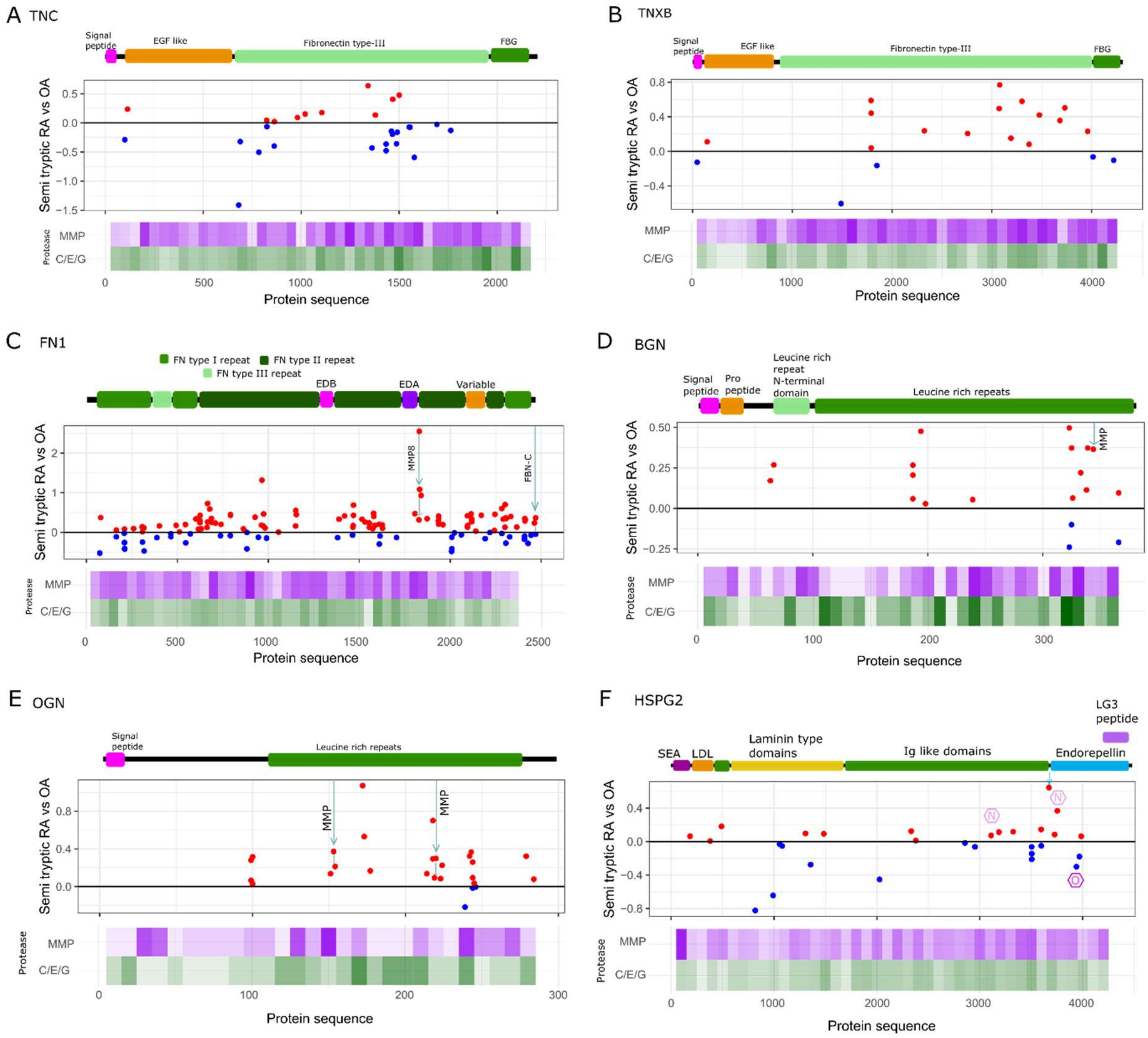


**Supplementary Fig. 4.**

(A) Cleavage map for TNC plotted as in Figure 3C. (B) for TNXB, (C) for FN1, (D) for BGN, (E) for OGN and (F) for HSPG2.

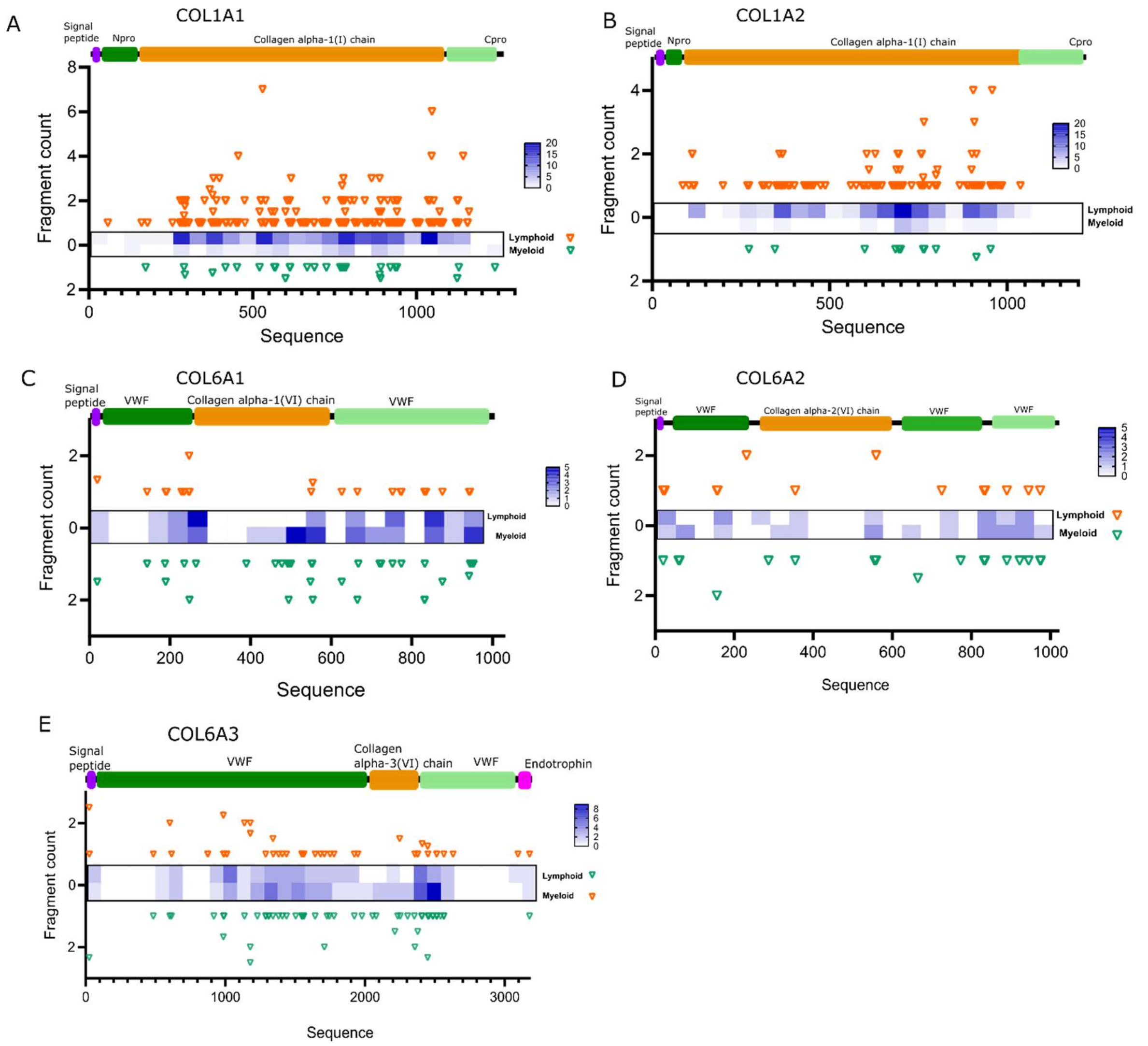


**Supplementary Fig. 5.**

Cleavage maps for (A) COL1A1, (B) COL1A2, (C) COL6A1, (D) COL6A2, (E) COL6A3. Orange and green triangles indicate fragments detected in lymphoid and myeloid samples respectively. Height represents PSMs for identified fragments in respective endotype. Heatmaps illustrate frequency of cleavage sites detected along the protein sequence.

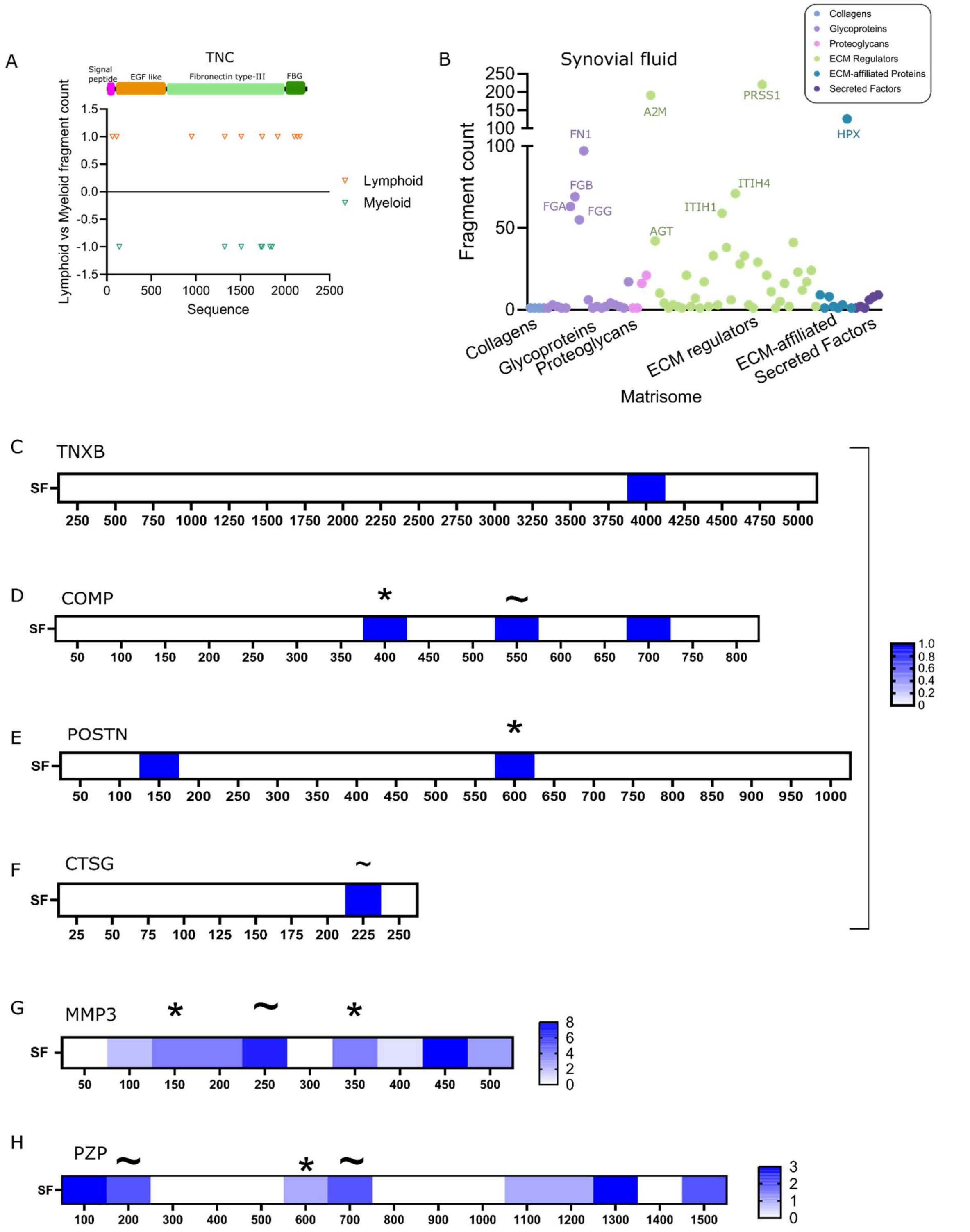


**Supplementary Fig. 6.**

(A) Cleavage map for TNC as in Figure 6A. (B) Total fragment counts for ECM proteins detected in synovial fluid from RA patients coloured by their matrisome group. The 10 proteins with the most fragments were labelled. (C) Heatmap of detected fragments from RA patient synovial fluid mapped along protein sequence for TNXB. (D) As in (C) for COMP. (E) As in (C) for POSTN. (F) As in (C) for CTSG. (G) As in (C) For MMP3. (H) As in (C) for PZP. For (C)-(H) Asterix mark fragments correlating with endotype-specific fingerprint. ~ mark fragments correlating with those previously detected. Proteins only shown if at least one semi-tryptic fragment was detected.
